# H105A peptide eye drops promote photoreceptor survival in murine and human models of retinal degeneration

**DOI:** 10.1101/2024.07.10.602890

**Authors:** Alexandra Bernardo-Colón, Andrea Bighinati, Shama Parween, Subrata Debnath, Ilaria Piano, Elisa Adani, Francesca Corsi, Claudia Gargini, Natalia Vergara, Valeria Marigo, S. Patricia Becerra

**Affiliations:** Section of Protein Structure and Function, Laboratory of Retinal Cell and Molecular Biology, National Eye Institute, National Institutes of Health; Bethesda, MD, USA; Department of Life Sciences, University of Modena and Reggio Emilia; 41125 Modena, Italy; CellSight Ocular Stem Cell and Regeneration Program, Sue Anschutz-Rodgers Eye Center, University of Colorado Anschutz Medical Campus; Aurora, Colorado, USA; Department of Pharmacy, University of Pisa; 56126 Pisa, Italy; Gates Center for Regenerative Medicine, Linda Crnic Institute for Down Syndrome and University of Colorado Alzheimer’s and Cognition Center, University of Colorado Anschutz Medical Campus; Aurora, Colorado, USA

## Abstract

Photoreceptor death causes blinding inheritable retinal diseases, such as retinitis pigmentosa (RP). As disease progression often outpaces therapeutic advances, finding effective treatments is urgent. This study focuses on developing a targeted approach by evaluating the efficacy of small peptides derived from pigment epithelium-derived factor (PEDF), known to restrict common cell death pathways associated with retinal diseases. Peptides with affinity for the PEDF receptor, PEDF-R, (17-mer and H105A) delivered via eye drops reached the retina, efficiently promoted photoreceptor survival, and improved retinal function in RP mouse models based on both the *rd10* mutation and the rhodopsin P23H mutation. Additionally, intravitreal delivery of AAV-H105A vectors delayed photoreceptor degeneration in the latter RP mouse model. Furthermore, peptide H105A specifically prevented photoreceptor death induced by oxidative stress, a contributing factor to RP progression, in human retinal organoids. This promising approach for peptide eye drop delivery holds significant potential as a therapeutic for preventing photoreceptor death in retinal disorders, offering a high safety profile, low invasiveness and multiple delivery options.

**One Sentence Summary:** Neurotrophic PEDF peptides delivered as eye drops preserve photoreceptor viability, morphology, and function in models of human retinal diseases.

## INTRODUCTION

Retinal degenerative diseases, such as inherited retinal dystrophies (IRDs), and age-related macular degeneration (AMD) are major causes of untreatable blindness. Retinitis pigmentosa (RP), the most common group of IRDs, is characterized by progressive photoreceptor cell loss that leads to blindness (*1, 2*). Although RP is a relatively rare genetic disorder, it is estimated to affect 1 in 4000 people worldwide (*3*). Its prevalence varies significantly depending on the population studied and the mutations involved in each RP type. Because the known RP genetic heterogeneity makes developing RP therapeutics rather challenging, gene-independent approaches (e.g., neuroprotective agents, retinal implants, gene therapy, dietary supplements, low-vision aids, and rehabilitation) aiming to treat a wider patient population have become more appealing (*2, 4–8*). Moreover, most ocular drug delivery routes for effective and specific targeting of photoreceptors (e.g., intravitreal or subretinal injections, intracameral injections using nanoparticles, liposome implants, intravenous administration, and others) involve various degrees of invasiveness (*9–11*). Importantly, as RP has a substantial impact on affected patients, finding potential RP treatments and identifying interventions to improve their quality of life are urgent clinical needs.

Current RP mouse models involve animals spontaneously mutated, or genetically engineered, replicating RP-associated genetic mutations in humans. Here, we selected two well-characterized models, *rd10* and *Rho^P23H/+^*mice that represent different genetic forms of RP with different rates of disease progression yet share photoreceptor cell death mechanisms. The *rd10* mouse is a model of autosomal recessive RP and has a spontaneous mutation in the *Pde6b* gene that inactivates phosphodiesterase PDE6, an enzyme involved in visual phototransduction (*12*). This mutation, also present in ∼5% human RP patients, leads to photoreceptor degeneration and simulates the disease process in humans (*1, 12–15*). We also used the *rd10/Serpinf1^−/−^*mouse model, in which pigment epithelium-derived factor (PEDF) deficiency increases retinal degeneration susceptibility (*16*). The Pro23His (P23H) is the most frequent mutation in the rhodopsin (*RHO*)-encoding gene, and alone accounts for ∼10% of autosomal dominant RP cases in North America. *Rho^P23H/+^* mice carry the P23H mutation engineered into the *Rho* gene and exhibit progressive retinal degeneration as in human patients (*17*). While in the *rd10* mouse photoreceptor death is completed in one month after birth, the photoreceptor degeneration in the *Rho^P23H/+^* mouse photoreceptor degeneration progresses much slower and most photoreceptors are absent by 180 days of age. Despite the different mutations, the mechanism of photoreceptor cell death in these two RP mutant retinas is associated with the calcium-calpain pathway. In the calcium-calpain pathway, an overload of intracellular calcium triggers calpain proteases, which then act on apoptosis-inducing factor through BAX activation (*18*). Furthermore, photoreceptor loss in RP models is also linked to oxidative stress (*18*). Thus, we used human stem cell-derived retinal organoids (ROs) undergoing oxidative stress-induced damage as they mimic hallmark features of retinal degeneration in RP (*19–21*).

The PEDF protein is known to prevent retinal degenerative processes by interfering with photoreceptor cell death pathways, is not toxic to humans or mice, and consequently holds significant promise as a useful RP therapeutic agent (*22–25*). One of the key benefits of PEDF is that its protective effects on photoreceptors are independent of the specific gene mutation causing the degeneration, thus offering a broad-spectrum approach to therapy. We have previously shown that PEDF activates the phospholipase activity of its receptor, PEDF-R, on retinal cells, which is crucial for its neurotrophic effects (*26–27*). Notably, ablation of PEDF-R leads to photoreceptor degeneration in mice, underscoring the importance of this pathway (*28*). In an effort to harness these protective effects, we have designed neuroprotective peptides from PEDF, namely 17-mer and its variant H105A (see **Fig. 1A**), which target PEDF-R and are involved in mitigating calcium overload in degenerating photoreceptors associated with RP (*29–33*). These short PEDF-derived peptides have potential benefits in neuroprotection (*22,23,29–33*), a key feature that makes them promising candidates for further research.

**Fig. 1:**
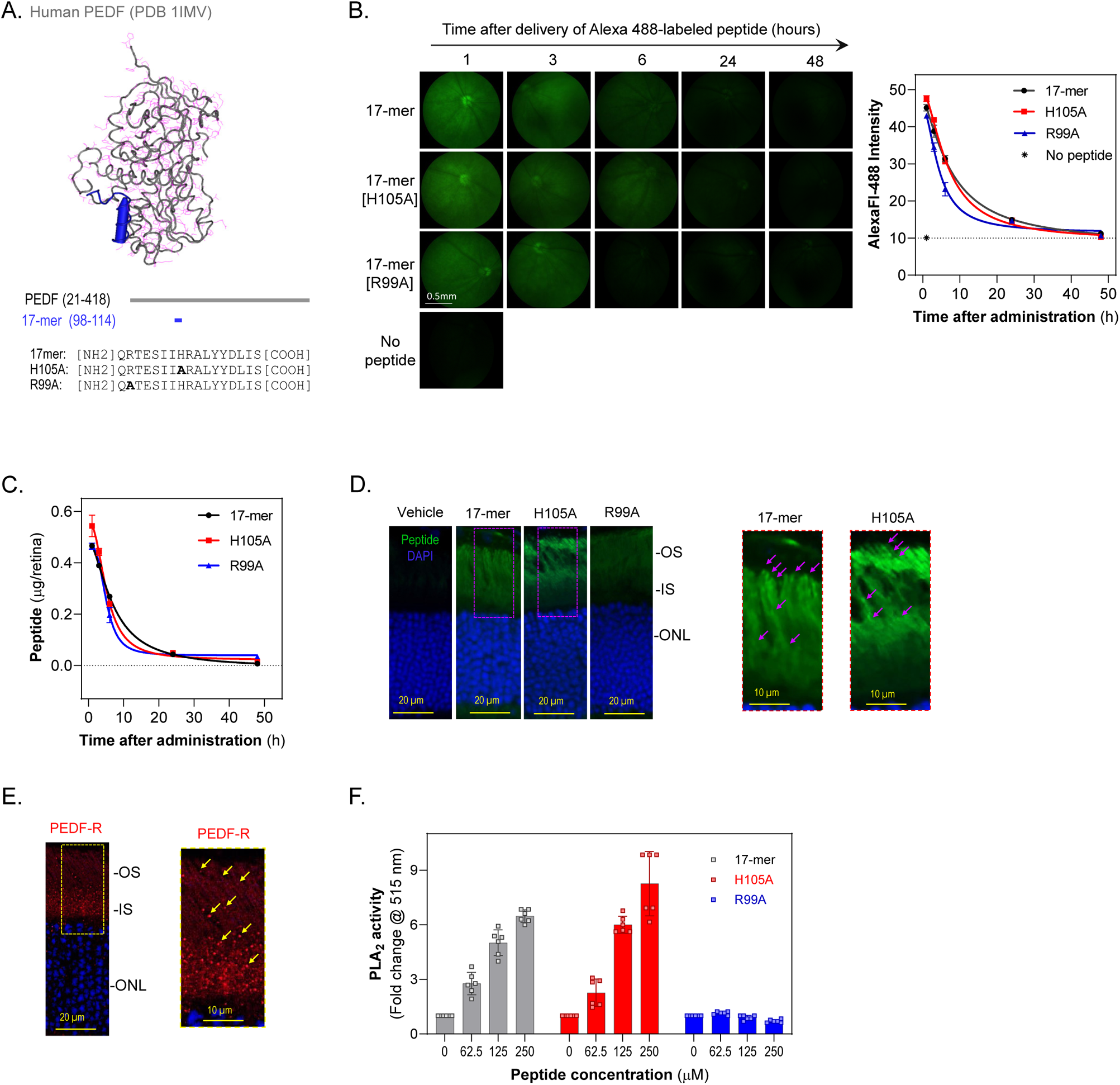
Penetration and Bioavailability of PEDF Peptides 17-mer, H105A, and R99A Delivered via Eye Drops in Mouse Retinas. (A) Peptide mapping and sequence. Tertiary structure of human PEDF highlighting the small peptide region with neurotrophic activity (17-mer) in blue (top). Primary structure of human PEDF showing the location of the 17-mer region (middle). Sequences of the 17-mer peptide and its variants, H105A and R99A (bottom). (B) In vivo detection of peptides. Fundus micrographs showing fluorescence from Alexa Fluor 488-labeled 17-mer, H105A, and R99A peptides in the eyes of C57Bl/6J mice at P21 days after eye drop administration of 5 µl of peptide at 1 mg/ml (left). Fundus images were taken at 1, 3, 6, 24, and 48 hours post-administration. Quantification of fluorescence intensity over time post-administration (right). The plot shows average intensity from 3 R.O.I.s per eye, with 5 eyes per time point. (C) Quantification of peptide in retinas. The plot shows the amounts of AlexaFluor-488-labeled peptides (17-mer, H105A, R99A) detected in dissected retinas at each time point. The amount of peptide was quantified from the fluorescence in retinal extracts using standard curves (Fig. S6A). Each data point represents the average ± S.D. from three eyes per time point. (D) Retinal cross-sections showing fluorescence distribution in the photoreceptor layer 1 hour after applying eye drops as in Panel B. Representative images for labeled 17-mer, H105A, and R99A peptides are shown, with magnified areas of 17-mer and H105A indicated by dotted rectangles. Magenta arrows point to punctuated areas in the outer segments (OS). IS, inner segments; ONL, outer nuclear layer. (E) Retinal sections showing PEDF-R distribution (red) and DAPI (blue) in C57Bl/6J mice at P21. Three retinas were used per group, with a representative image shown. A magnified area (dotted rectangle) is provided. (F) Effects of 17-mer, H105A, and R99A peptides on the PLA_2_ activity of PEDF-R. Peptides were preincubated with recombinant human PEDF-R[1-288] for 30 min, and PLA_2_ activity was measured. Each bar represents the average fold-change in activity over control (no peptide), from six replicate enzymatic reactions with indicated peptide concentrations and final concentration for PEDF-R of 60 nM.

The delivery method of these peptides is a critical factor for their efficacy. This study explores whether administering PEDF peptides via eye drops -an understudied delivery method for targeting photoreceptors-, could effectively deliver these peptides to the retina. The goal is to determine whether these delivery methods can delay, stabilize, or prevent progression of IRDs such as RP. To address these questions, our research investigates the survival effects of PEDF peptides on photoreceptors using preclinical mouse models of RP. Additionally, we explore an alternative delivery method involving gene therapy vectors (currently FDA-approved for other indications) as well as testing in human ROs. These models help assess the potential therapeutic benefits and practicality of PEDF peptide delivery methods, paving the way for future translational research and potential clinical applications in RP treatment.

## RESULTS

### Small PEDF peptides delivered via eye drops penetrate into mouse retinas

Previously, we demonstrated that neurotrophic PEDF-derived peptides 17-mer and H105A have affinity for PEDF-R, and that R99A peptide lacks PEDF-R affinity and neurotrophic activity (*29*). To evaluate their retinal penetration upon delivery via eye drops, solutions of Alexa Fluor™ 488-conjugated peptides were applied as eye drops to C57Bl/6J mice. Peptide penetration into the photoreceptor layer was monitored by fluorescence fundoscopy at time intervals up to 48 hours post-administration. After 1-hour, fundi images exhibited high fluorescence intensity, which diminished over time **(Fig. 1B).** About 10% of applied peptide amounts reached the posterior retina within one hour, with 5% −1% remaining between 6 – 24 hours, respectively, and becoming undetectable by 48 hours for all three peptides (**Fig. 1C**). Peptide distribution in the retina was specific to photoreceptor inner (IS) and outer segments (OS) at 1-hour (**Fig. 1D**) and 6-hours post-administration, increasing the intensities in proportion with their PEDF-R affinity R99A<17-mer<H105A (*29*). While R99A was diffusely distributed, peptides 17-mer and H105A decorated the OS in a punctuated fashion (**Fig. 1D**) that matched the PEDF-R distribution in the retina (**Fig. 1E**). No visible pathologic changes were noticed in retinas exposed to the PEDF peptides. Because the phospholipase A2 (PLA_2_) activity of PEDF-R is critical for the PEDF-mediated photoreceptor protection (*27*), we determined the PLA_2_ activity in response to the PEDF peptides. While the presence of additional PLA_2_ enzymes in the mouse retina precluded evaluating the peptides for stimulating PEDF-R in this tissue, assessments using purified recombinant PEDF-R in the presence of peptides showed that only 17-mer and H105A peptides are functional effectors of PEDF-R capable of stimulating its PLA_2_ activity (**Fig. 1F**).

These findings imply that PEDF-R affinity facilitated the localization of peptides at photoreceptors, highlighting the effective bioavailability of 17-mer and H105A peptides delivered as eyedrops to activate PEDF-R in the photoreceptor layer. Additionally, the observed clearance of peptides within 24 hours suggested that daily administration of peptide eye drops might ensure prolonged bioavailability, thereby potentially enhancing their therapeutic efficacy.

### Eye drops of 17-mer and H105A peptide prevent photoreceptor death in *rd10* mouse models of RP

Phosphatidyl serine (PS) externalization by phospholipid translocases provides an emblematic eat-me signal in apoptotic cells for efferocytosis. This has been exploited for detecting rat photoreceptor cell death *in vivo* using the PSVue®-550 fluorescent probe, which contains a bis(zinc2^+^dipicolylamine) (Zn-DPA) group that selectively binds to cell membranes enriched with exposed PS phospholipids in apoptotic and necrotic cells (*33,34*). We optimized this method to monitor PS externalization in photoreceptors of *rd10* and *rd10/Serpinf1^−/−^* mice starting at P16, when their retinas start showing histological changes consistent with retinal degeneration (*13*), until P25 *in vivo,* when most of the photoreceptors have died (**Fig. S1**). We found that PS externalization agreed with the natural history of photoreceptor death, and selected P21 as the endpoint for PS externalization monitoring *in vivo*.

To evaluate the efficacy of eye drops containing 17-mer, H105A and R99A peptides in delaying PS externalization in both *rd10* and *rd10/Serpinf1^−/−^* mouse models *in vivo*, animals received daily eye drops starting at P15 before photoreceptor death onset. Treatment continued until P20, when PSVue®-550 was also applied, and fundoscopy was performed at P21 (**Fig. 2A**). The results demonstrated that eye drops containing 17-mer or H105A diminished fundi fluorescence, indicating lower PS externalization compared to the vehicle (HBSS) eye drops. Notably H105A peptide was more effective than 17-mer, while R99A peptide showed no significant efficacy (**Fig. 2B**). Further titration experiments with 17-mer and H105A peptides revealed concentrations as low as 0.5 mg/ml were effective in restraining PS externalization in photoreceptors in both RP mouse models (**Fig. 2C**). This demonstrates the benefit of both 17-mer or H105A peptide eye drops in blocking PS externalization, an early step in the cell death process. For subsequent experiments, a working dose of 1 mg/ml peptide was used.

**Fig. 2:**
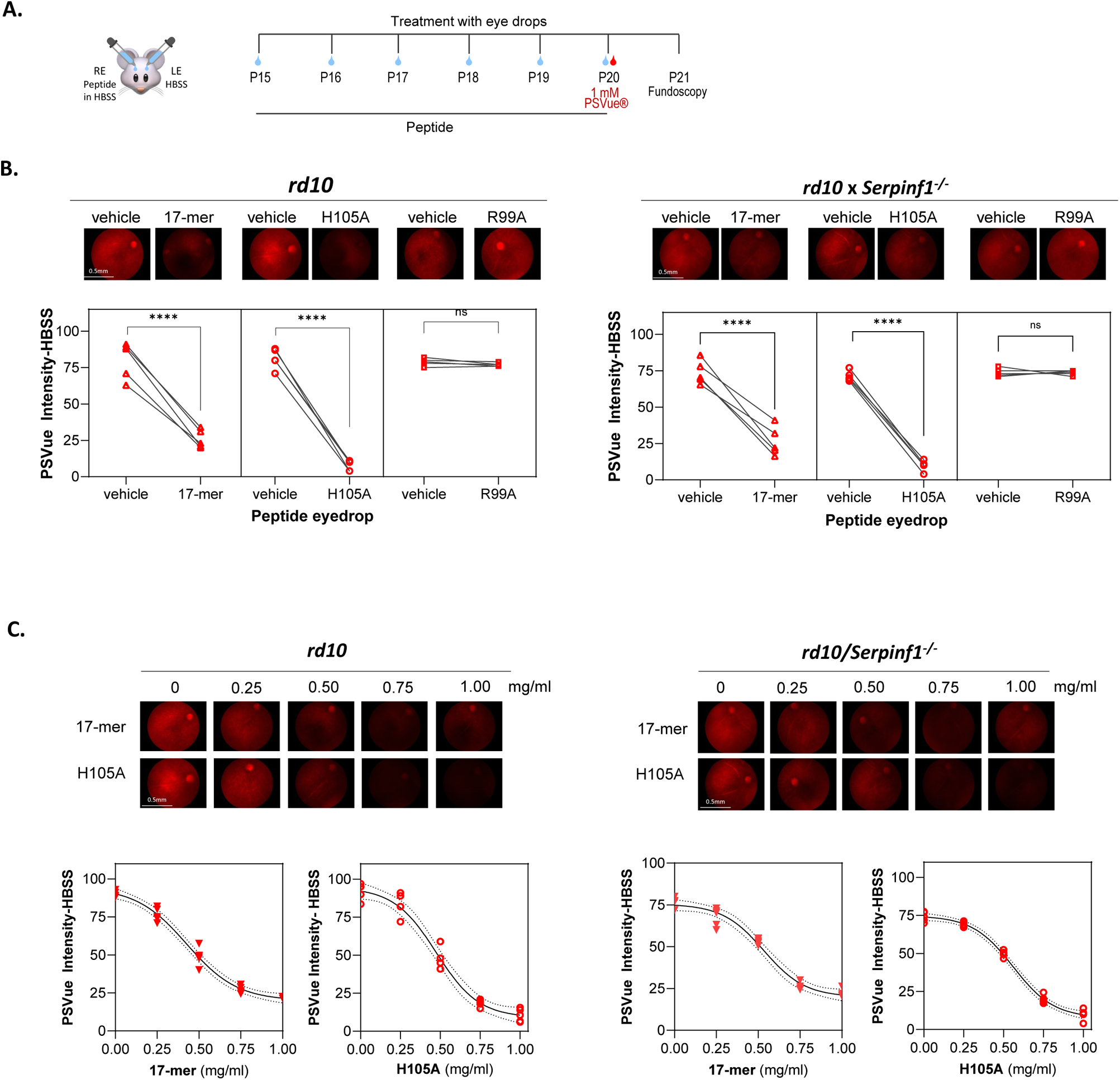
Effect of PEDF peptide eye drops on protection against photoreceptor cell death in *rd10* and *rd10/Serpinf1^−/−^* mice. (A) Illustration of the experimental timeline of peptide application. Eye drops containing 5 µl of 17-mer, H105A, or R99A peptides in HBSS were administered daily to eyes of *rd10* and *rd10/Serpinf1^−/−^* mice from P15 to P20. Vehicle (HBSS) was applied in contralateral eyes. At P20, PSVue® was applied via eye drops, and fluorescence fundoscopy was conducted at P21 to assess photoreceptor cell death. RE and LE refer to the right and left eyes, respectively. (B) Fluorescence fundoscopy micrographs. Representative fluorescence fundoscopy images of retinas from mice treated as in Panel A with eyedrops of 1 mg/ml of each peptide. Quantification of fluorescence intensity was performed using ImageJ, with average intensity values corrected for background fluorescence from eyes that had not received PSVue or peptides. Each line represents data from one animal comparing treated and contralateral eyes, with each data point being the average of three R.O.I.s per eye, and with 5 eyes per condition. Statistical significance was determined by unpaired t-test (****p < 0.0001). Scale bar = 0.5 mm. (C) Dose response of peptide eye drops. Representative fluorescence fundoscopy micrographs showing retinas treated with varying concentrations of 17-mer and H105A peptides (0.0 mg/ml to 1.0 mg/ml) in *rd10* and *rd10/Serpinf1^−/−^*mice. Fluorescence intensity was quantified and analyzed using Sigmoidal interpolation in GraphPad. Each data point represents the average of 3 images per retina, with a total of 5 retinas per condition.

Given that intravitreal injections of PEDF protein, 17-mer or H105A peptides elevate anti-apoptotic BCL-2 and attenuate pro-apoptotic BAX levels in photoreceptors of retinal degeneration models such as the *rd1* mouse and RCS rat (*30,36*), we hypothesized that these peptides delivered via eye drops would similarly regulate BCL-2 and BAX proteins to prevent cell death in *rd10* and *rd10/Serpinf1^−/−^*retinas. To test this hypothesis, we assessed the distribution of BAX and BCL2 proteins in photoreceptors of *rd10* and *rd10/Serpinf1^−/−^* mice following the application of peptide eye drops. Our results showed that both 17-mer and H105A eye drops significantly decreased BAX and increased BCL2 levels in the photoreceptors in both RP models (**Figs. 3A, 3B)**.

**Fig. 3:**
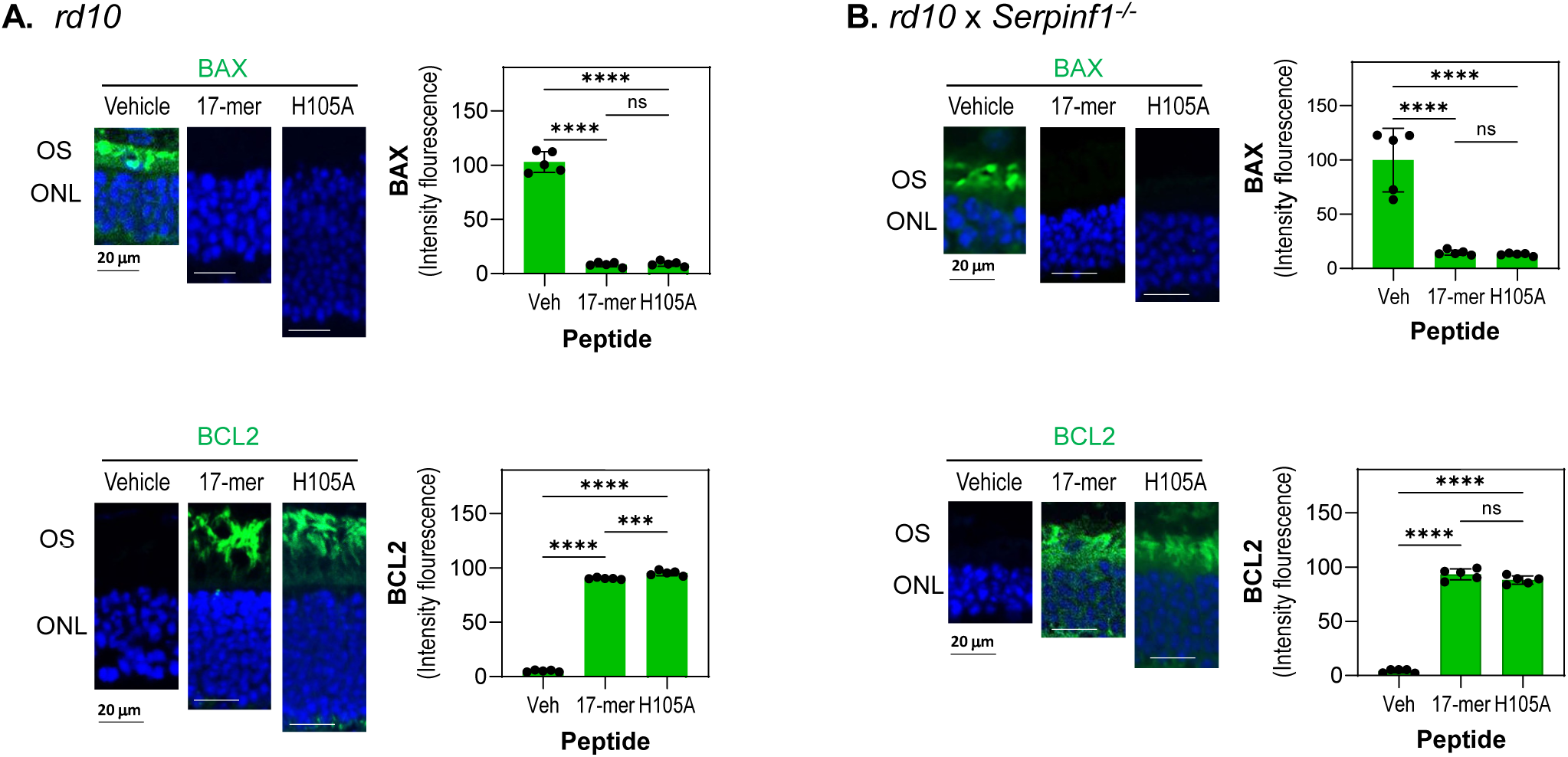
Eyedrops of 17-mer and H105A peptides decreased the BAX/BCL2 ratio in photoreceptors of rd10 and rd10/Serpinf1-/-mice. (A and B) Immunostaining of retinas. Representative fluorescent micrographs of retinas from *rd10* (A) and *rd10/Serpinf1^−/−^* (B) mice treated with 17-mer or H105A peptides at 1 mg/ml via eye drops, along with vehicle-treated controls. Immunostaining was performed with antibodies against BAX or BCL2 (green), and DAPI (blue). Histograms show quantification of immunofluorescence intensity for BAX and BCL2 markers. Five retinas per group were analyzed, with two sections per retina. Each individual data point represents the average of one retina, with statistical significance determined by unpaired t-test (***p < 0.0001, ****p < 0.00001). Scale bar = 40 µm.

Together, the findings indicate that daily treatments with 17-mer or H105A peptide eye drops effectively protected *rd10* and *rd10/Serpinf1^−/−^* mice against photoreceptor cell death, demonstrating the potential of these peptides as a non-invasive approach for RP.

### Eye drops of 17-mer and H105A peptides improve photoreceptor morphology and function in *rd10* mouse models

Early changes effected by RP can be histologically detected as shortening of photoreceptor OS and photoreceptor loss. Histological examination showed that retinas of *rd10* and *rd10/Serpinf1^−/−^* mice treated with daily H105A or 17-mer peptide eye drops had longer OS and thicker outer nuclear layer (ONL, containing photoreceptor cells) than vehicle-treated or untreated eyes, while R99A had no effect at P21 (**Figs. 4A, 4B**). In *rd10* mice, H105A or 17-mer peptide eye drops restored ONL thickness, with H105A to almost wild-type levels (**Figs. 4A, 4B**; spider plots). In *rd10/Serpinf1^−/−^*, ONL preservation was less marked, likely due to the increased retinal degeneration susceptibility caused by *Serpinf1* gene deletion (*16*). Again, vehicle- and R99A-treated eyes were inefficient in restoring ONL thickness. Application of H105A eye drops every other day and extending the endpoint to P25, when most photoreceptors degenerate, also improved *rd10* photoreceptor morphology albeit with lower efficiency (**Fig. S2**).

**Fig. 4:**
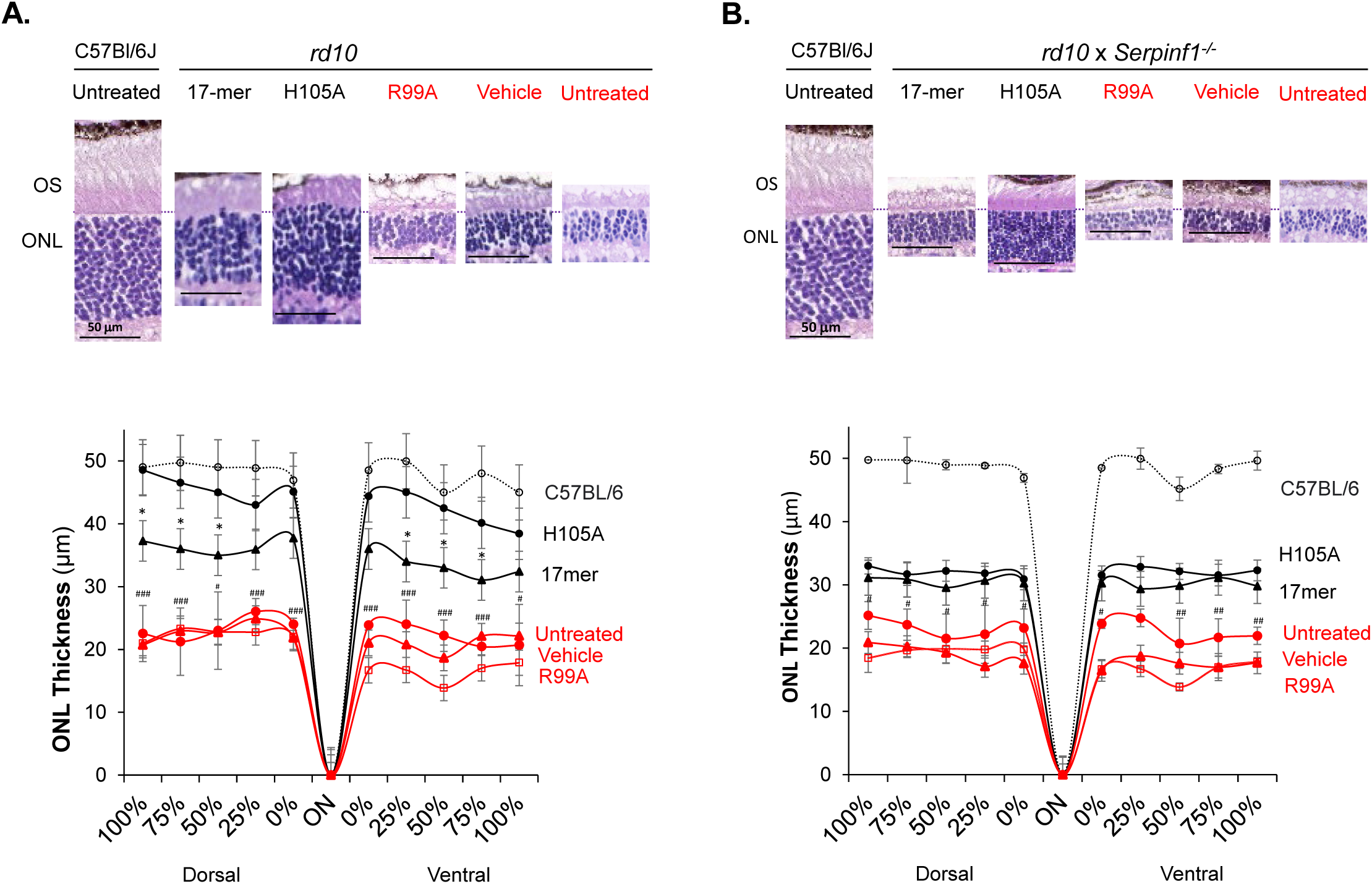
Effect of eyedrops of 17-mer and H105A peptides on photoreceptor morphology of rd10 and rd10/Serpinf1-/-mice. (A and B) Histological Evaluation. Representative microphotographs of retina sections from rd10 (A) and rd10/Serpinf1-/- (B) mice treated with 17-mer, H105A, or R99A peptides at 1 mg/ml, or with vehicle (HBSS), compared to wild type C57Bl/6J mice. Sections were stained with hematoxylin and eosin. Spider Plot illustrate the thickness of the outer nuclear layer (ONL) in rd10 (C) and rd10/Serpinf1-/- (D) mice. Five retinas per group were analyzed and each data point represents the average ± SEM per location relative to the optic nerve (ON), analyzed using One-way ANOVA. Statistics for comparisons between H105A and vehicle is indicated by #p < 0.05, #p < 0.05, ##p < 0.01, ###p < 0.001). Statistics for comparisons between H105A and 17-mer is indicated by *p < 0.05. Scale bar = 50 µm.

RP is associated with reduced or delayed electrical responses to light. Electroretinography (ERG), the standard technique for assessing retinal dysfunction associated with RP and for measuring RP treatment outcomes, was performed to evaluate the functional effects of 17-mer or H105A eye drops in *rd10* and *rd10/Serpinf1^−/−^* mice. ERG a-wave and b-wave responses are indicative of photoreceptor function, and of post-synaptic activation of bipolar cells by photoreceptors, respectively. In dark-adapted *rd10* and *rd10/Serpinf1^−/−^* mice at P21, the peptides elicited a- and b-wave responses that increased exponentially with light stimuli between log(−0.01) and log(10) cd.s/m^2^, indicating that, relative to vehicle-treated eyes, the PEDF-derived peptides protected visual function (**Fig. 5)**. Application of H105A peptide eye drops every other day and extending the endpoint to P25 also improved the function of *rd10* photoreceptors, with slightly lower efficacy than daily treatments and a shorter endpoint (**Fig. S2**).

**Fig. 5:**
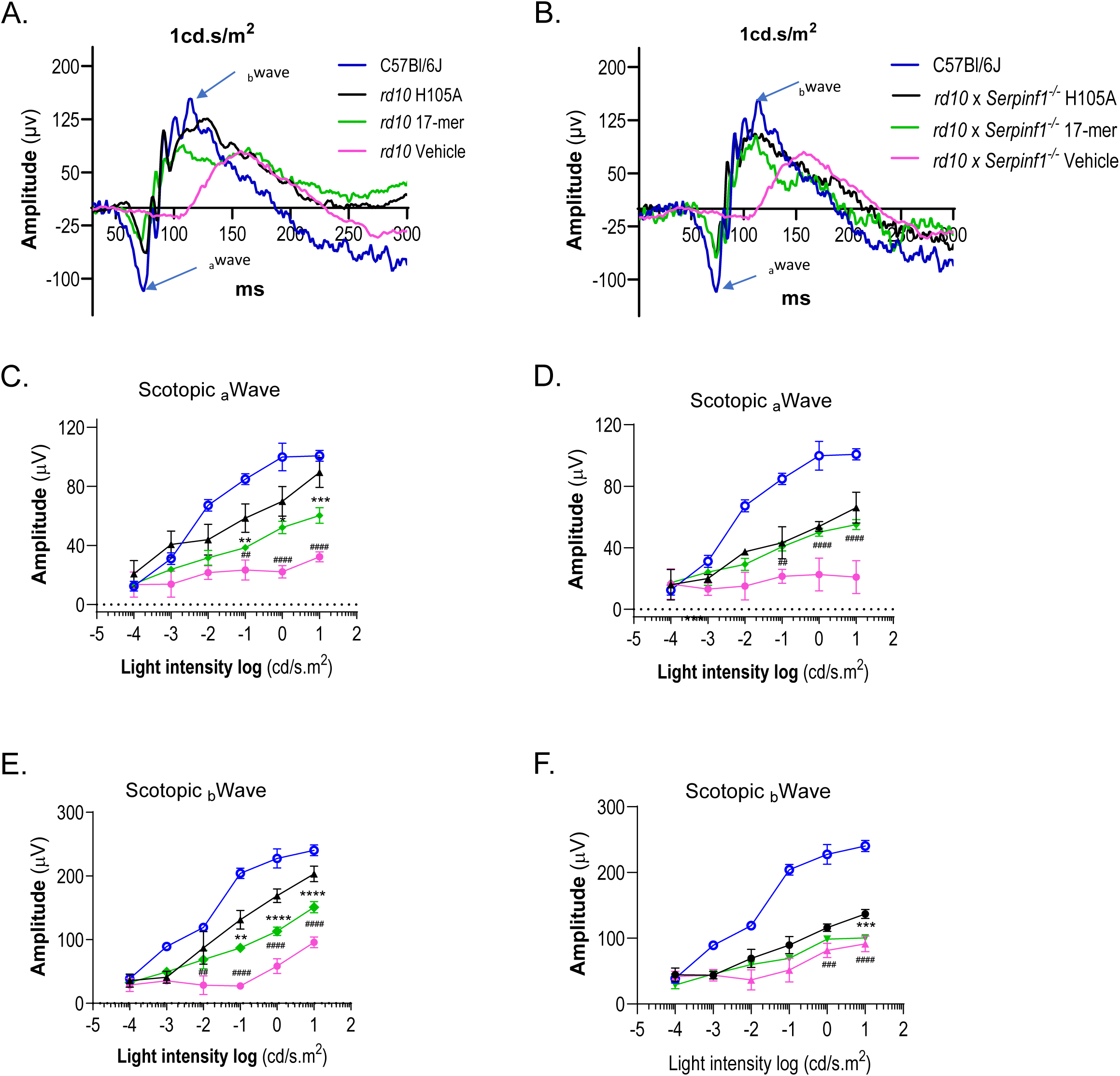
Effect of daily eyedrops of PEDF peptides on ERG a-wave and b-wave retinal function. (A and B) Representative ERG waveforms for *rd10* (A) and *rd10/Serpinf1^−/−^* (B) mice at P21 in response to a light flash (1 cd.s/m²). Amplitude (y-axis) is plotted against time in milliseconds (ms, x-axis). Treatments are indicated to the right of each plot and are color coded, with control wild type C57Bl/6J mice without peptide treatment shown in blue. A-wave and b-wave are indicated by arrows. (C and D) Photoreceptor a-wave amplitudes. Averages of photoreceptor a-wave amplitudes for *rd10* (C) and *rd10/Serpinf1^−/−^* (D) mice as a function of light intensity (cd/s.m², x-axis). Responses of animals treated with 17-mer are shown in green, H105A in black, and vehicle (HBSS) in red. Wildtype responses are shown in blue. (E and F) Bipolar cell b-wave amplitudes. Averages of bipolar cell b-wave amplitudes for *rd10* (E) and *rd10/Serpinf1^−/−^* (F) mice as a function of light intensity (cd/s.m², x-axis), with treatments indicated as in panels (C) and (D). The number of mice evaluated for the data shown in Panels C-F were n = 8 for each *rd10*, *rd10/Serpinf1*^−/−^ and wild type. Each data point represents the average ± SD of each genotype, analyzed by unpaired t-test. Statistical significance between H105A and vehicle is indicated by ##p < 0.01, ###p < 0.001, ####p < 0.0001, and between H105A and 17-mer by **p < 0.01, ***p < 0.001, ****p < 0.0001.

These findings show the ability of H105A or 17-mer peptide eye drops to limit photoreceptor cell morphology changes and loss, to improve electrical light responses in both mouse models, and thus delay retinal degeneration.

### H105A peptide eye drops activate photoreceptor survival and improve morphology of the ONL in the *Rho^P23H/+^* mouse model of RP

We evaluated the efficacy of eye drops containing the H105A peptide in preventing photoreceptor cell death in the *Rho^P23H/+^* mouse model of RP, which exhibits a slower progression of photoreceptor degeneration compared to *rd10* mice. Daily administration of H105A peptide eye drops from P14, before the onset of photoreceptor death in *Rho^P23H/+^* mice (*37*), to P19 was performed with contralateral control eyes treated with vehicle HBSS. (**Fig. 6A**). Peptide H105A attenuated the levels of BAX and increased BCL2, indicating the activation of survival pathways and restraint of apoptotic factors (**Fig. 6B**). Assessment of ONL thickness at P19, labeled by rhodopsin, revealed significant preservation in the ventral part of the retina, which degenerates faster in this RP mouse model (*17*) (**Fig. 6C**). Quantification of ONL thickness, represented in a spider graph, confirmed significant preservation in the ventral retina (**Fig. 6D**). The outer segments (OS) of rod photoreceptors, labeled by rhodopsin, were significantly preserved with H105A peptide treatment (**Fig. 6E**). These results demonstrate that H105A peptide eye drops effectively promote photoreceptor survival and preserve retinal structure in the *Rho^P23H/+^*mouse model of RP, particularly in regions of the retina that are more susceptible to degeneration. This suggests a promising therapeutic approach for treating retinal degenerative diseases.

**Fig. 6:**
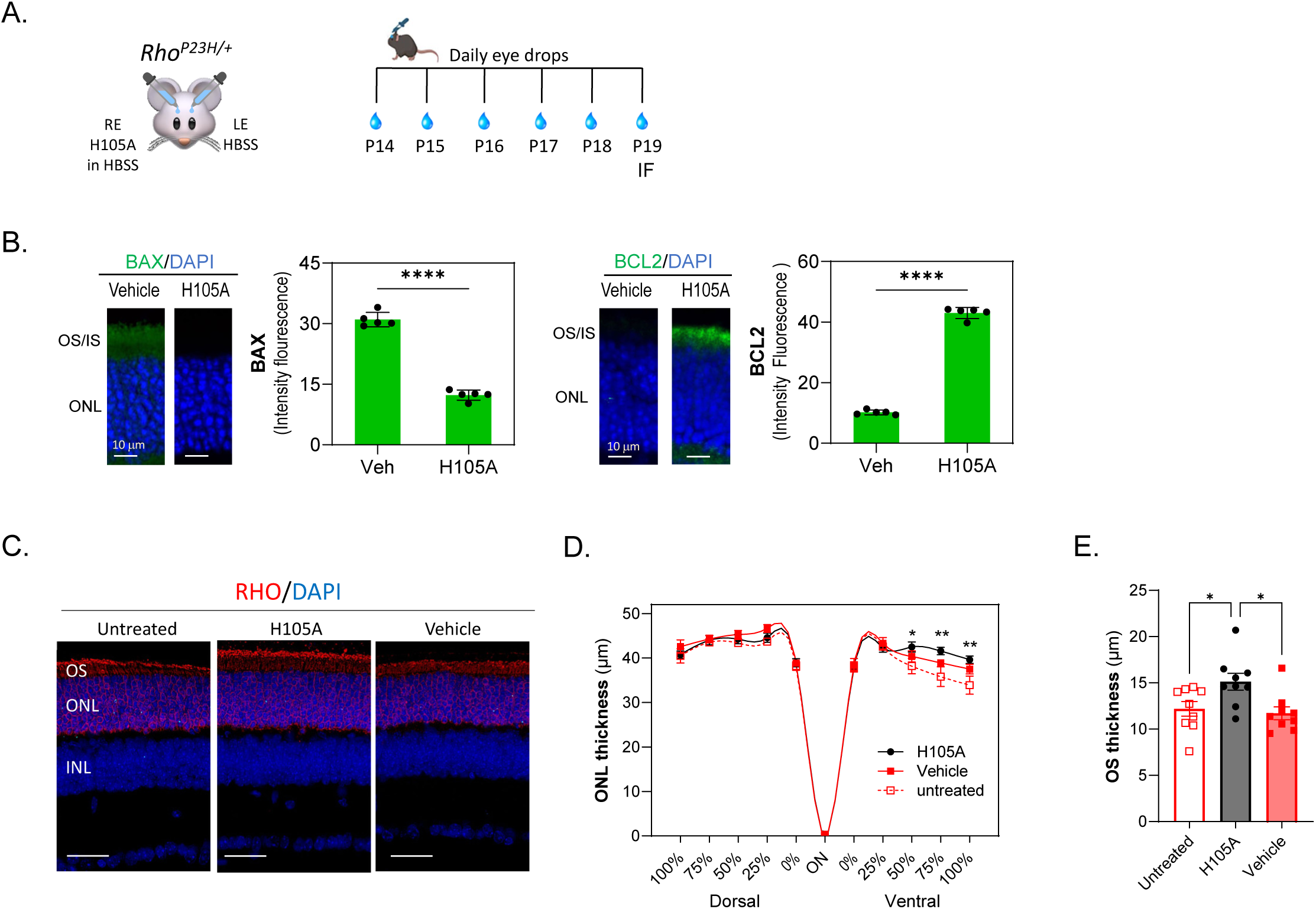
Effects of eye drops containing H105A peptide in the photoreceptors of RhoP23H/+ mice. (A) Scheme of experimental design. Daily peptide eye drops (5 µl of 1mg/ml peptide) were applied to mice starting at P14 until P19 when immunofluorescence of retinal cross sections was performed. The contralateral control eyes were treated with vehicle HBSS. (B) Immunostaining was performed with antibodies against BAX or BCL2 (green), and DAPI (blue). Histograms show quantification of immunofluorescence intensity for BAX and BCL2 markers. Five retinas per group were analyzed, with two sections per retina. Each individual data point represents the average of one retina, with statistical significance determined by unpaired t-test (****p < 0.00001). Scale bar = 10 µm. (C) Assessment of ONL and OS at P19, labelled by rhodopsin (red), revealed a significant preservation upon H105 peptide exposure. (D) Quantification of ONL thickness represented in the spider graph confirmed a significant preservation in the ventral retina. Each data point corresponds to the average ± SEM, with statistical analysis performed using Two-way ANOVA with Šídák’s multiple comparisons for each group: ∗ p<0.05; ∗∗ p<0.01 (E) Quantification of OS thickness labelled by rhodopsin shows preservation of photoreceptor OS with H105A peptide treatments. n = 9 for each condition. Presented as average ± SEM, with statistics performed using One-way ANOVA ∗ p<0.05

### Delivery of H105A by AAV transduction protects photoreceptors in the *Rho^P23H/+^* mouse model of RP

To assess the longer-term effects of H105A peptide, we developed a gene therapy-based delivery system for sustained expression of H105A. We selected the adeno-associated virus serotype 2 (AAV2) due to its efficacy in transducing retinal cells, making it well-suited for treating ocular disorders (*38*). AAV2 is a well-established and FDA-approved viral vector. Since the therapeutic agent is secreted extracellularly and does not require direct expression by photoreceptors, we opted for intravitreal (IVT) injection. This method is safer than subretinal delivery as it avoids the risk of retinal detachment. Additionally, IVT injection targets cells across the entire retina providing a broader area of treatment compared to the localized effect of subretinal injections.

A single IVT of AAV-H105A was administered to the eyes of mice at P5, with a control AAV-GFP IVT injection in the contralateral eye (**Fig. 7A**). By P19 and P180, cells transduced with AAV-H105A expressed viral mRNA and produced the neuroprotective agent in the retina (**Figs. 7B, S3**). At P19, AAV-H105A significantly decreased BAX levels and increased the BCL2 levels **(Fig. 7C)**. Additionally, it reduced the percentage of TUNEL-positive photoreceptor nuclei compared to GFP controls (**Fig. 7D**), indicating that AAV-H105A IVT protected the *Rho^P23H/+^*retinas against photoreceptor cell loss. Retinal assessments at P19 and P180 revealed slightly thicker ONL and better preservation of rhodopsin and PNA staining in the OS, which label rods and cones, respectively, in eyes producing H105A compared to GFP (**Figs. 7E, 7F**). Functional evaluations using ERG demonstrated improved b-wave responses in both scotopic (rods) and photopic (cones) conditions, with significant improvement in rod function and slight improvement in cone function six months after AAV delivery (**Figs. 7G, 7H**).

**Fig. 7:**
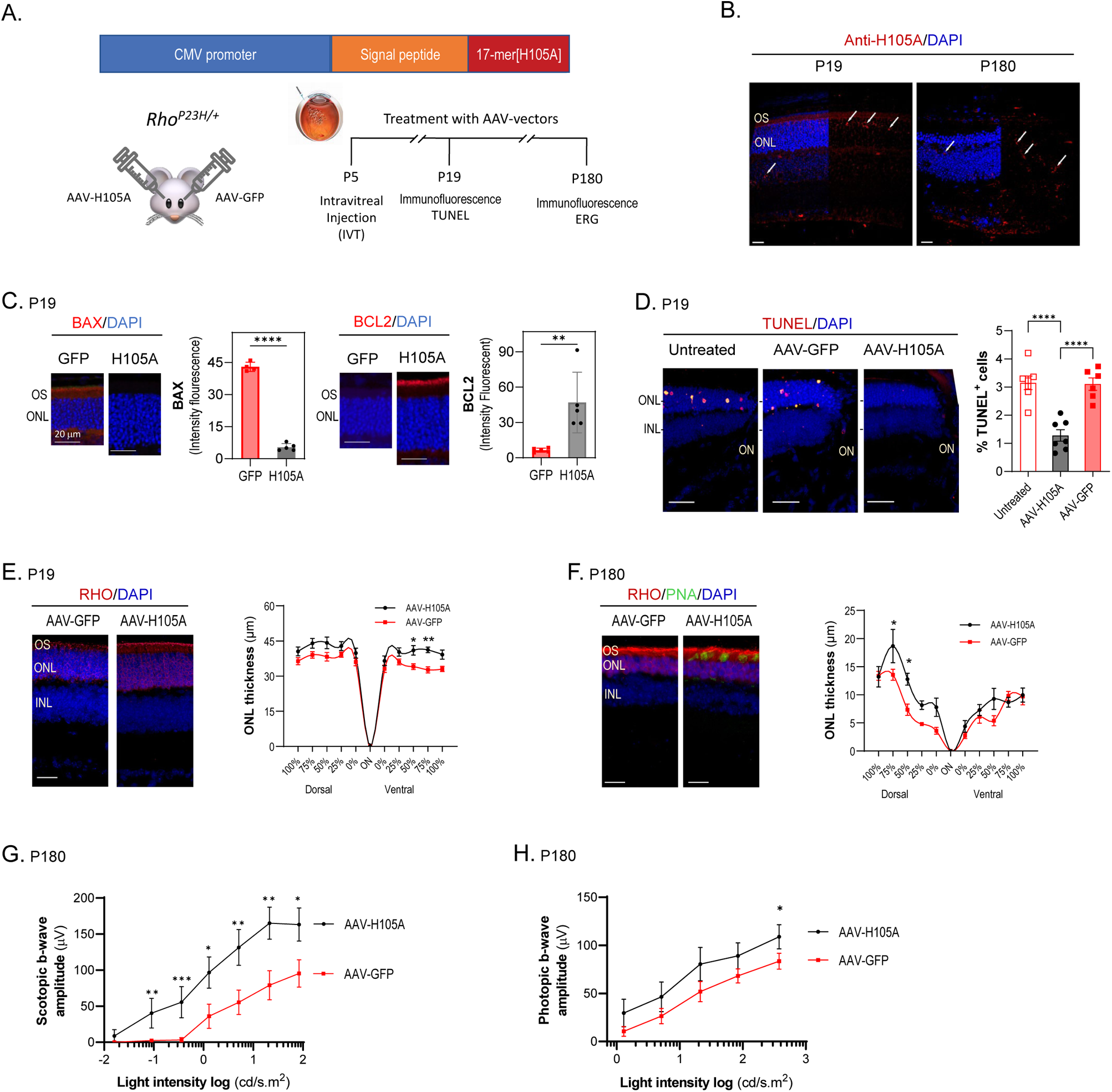
Effect of AAV-H105A intravitreal injection on the *Rho^P23H/+^* mouse model of RP. (A) Scheme for the experimental design illustrating the cDNA construct containing the secretion signal peptide of interferon beta in frame with the H105A coding sequence under the control of the CMV promoter in the AAV vector. One single IVT injection of 0.5 µl of AAV-H105A (1.9 × 10^12^ GC/ml) or AAV-GFP (4.5 × 10^12^ GC/ml) (as control) was administered in eyes of *Rho^P23H/+^* mice at P5. Assessments were performed at P19 and P180 by immunofluorescence, and TUNEL of retina sections, and at P180 ERG in live animals. (B) Immunostaining of retinal cross-sections of *Rho^P23H/+^* mice at P19 and P180 treated with AAV-vectors as described in Panel A, showing specific detection of H105A peptide (red), with nuclei stained with DAPI (blue). White arrows indicate H105A detection in different retinal layers. Scale Bar: 20 µm. (C) Immunostaining of retinal cross-sections of mice at P19, treated as described in Panel A, with antibodies to BAX or BCL2 (red), as indicated, and with DAPI (blue). Histograms show quantification of the fluorescence signal. Five retinas per group were analyzed, with two sections per retina. Each data point corresponds to the average ± SD, with statistical analysis performed using One-way ANOVA for each group: ∗∗ p<0.01; ∗∗∗∗ p<0.0001. Scale Bar: 20 µm. (D) TUNEL of retinal cross-sections of mice at P19 treated as described in Panel A, showing TUNEL-positive cells (red) with DAPI (blue). Histograms show quantification of the percentage of TUNEL+ photoreceptors. N = 8 per condition. One-way ANOVA with Šídák’s multiple comparisons for each group ****p < 0.0001. Scale Bar: 20 µm. ON, optic nerve. (E) and (F) ONL thickness in retinas treated as described in Panel A, assessed at P19 (E) and P180 (F), with retinal cross-sections stained with DAPI (blue) and anti-rhodopsin (red). Plots for ONL thickness as a function of distance from the optic nerve (ON) are shown. For ONL thickness, Two-way ANOVA with Šídák’s multiple comparisons for each group was used. For OS thickness, unpaired t-test was used ∗ p<0.05; ∗∗ p<0.01. Scale Bar: 20 µm. (G) and (H) Visual function assessment by ERG recording at P180 under scotopic (G) and photopic (H) conditions. Sensitivity curves in terms of b-wave amplitude in response to different light stimuli are shown. Each data point corresponds to the average ± SEM, with statistical analysis performed using One-way ANOVA for each group: ∗ p<0.05; ∗∗ p<0.01; ∗∗∗ p<0.001.

These results underscore the multifaceted benefits of H105A peptide delivery, including enhanced photoreceptor survival, improved morphology, and augmented light responses in the *Rho^P23H/+^* model of RP, especially considering the model’s slow photoreceptor degeneration progression.

### PEDF-derived H105A peptide promotes retinal cell survival in hiPSC-derived human retinal organoids

To further demonstrate the efficacy of the H105A peptide in promoting retinal photoreceptor and neuronal survival, we used human induced pluripotent stem cell (hiPSC)-derived retinal organoid (RO) technology. RO models mimic the native retinal three-dimensional structure and cellular makeup, including photoreceptors with inner and outer segments, and light responsiveness (*39*). To generate a disease-relevant paradigm of photoreceptor cell death, we cultured human ROs up to 180 days of differentiation (D180), a time at which the ONL is well developed, and all photoreceptor subtypes have been specified and are relatively mature (*39,40*) and treated them with cigarette smoke extract (CSE), a potent oxidant commonly used to induce macular degeneration-related damage in retinal cultures (*40–43*). Although RP is a hereditary disease, non-genetic factors, such as oxidative stress, play a central role in its pathogenesis and progression (*18–21*). ROs were simultaneously treated with CSE and with H105A or the R99A control peptide (**Figs. 8A, 8B**), using CSE alone or vehicle (DMSO) as controls.

**Fig. 8:**
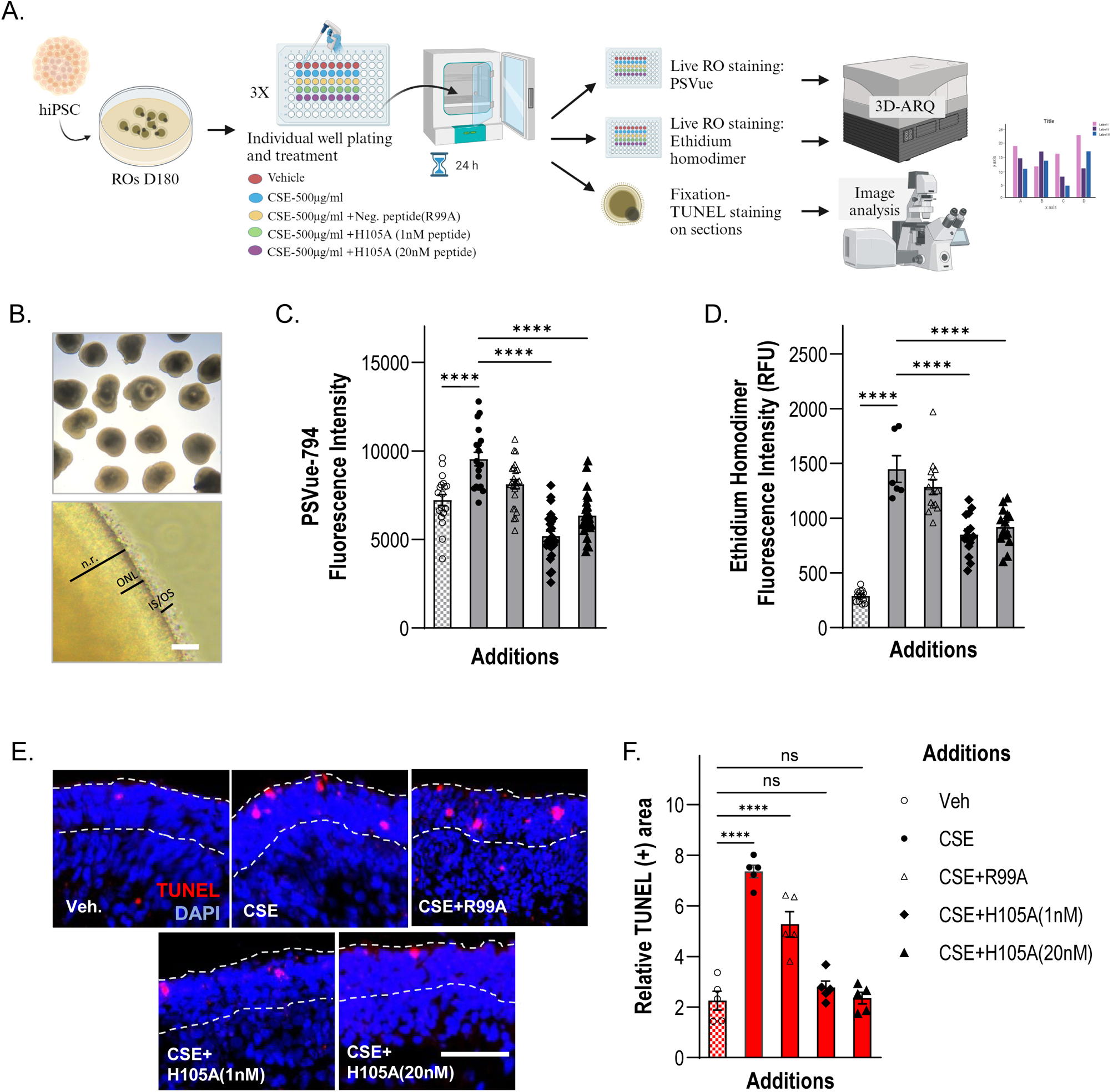
Effect of H105A peptide on CSE-induced retinal cell death in human iPSC-derived retinal organoids. (A) Diagram of Experimental Design. Human ROs at 180 days of differentiation (D180) were treated with vehicle (DMSO), CSE(500μg/ml), CSE+R99A(20nM), CSE+H105A(1nM), or CSE+H105A(20nM) for 24 h, and cell death was assessed by PSVue794 or Ethidium homodimer staining of live organoids or by TUNEL staining in fixed RO sections. Diagram created with BioRender.com (B) Representative image of retinal organoids. Top panel shows a representative bright field image of ROs at D180. Bottom panel is a close-up of a D180 RO showing neural retina (n.r.), outer nuclear layer (ONL), and inner and outer segment structures of photoreceptors (IS/OS). Scale Bar: 50 μm. (C) PSVue794 fluorescence intensity (RFU) of PSVue794 in live ROs under the 5 experimental conditions. CSE increases early apoptotic events, evidenced by a statistically significant increase in PSVue794 intensity compared with Vehicle control, and H105A prevents this increase at both tested concentrations. CSE+H105A(1nM) = 5185.2 ± 275.4 and CSE+H105A(20nM) = 6342.7 ± 261.3, vs. CSE alone = 10172.6 ± 431.0 and CSE+R99A(20nM) = 8112.5 ± 266.2. Mean ± SEM; **** p<0.0001, One-way ANOVA. (D) Fluorescence intensity (RFU) of Ethidium homodimer in live ROs under each treatment condition. CSE increases cell death, evidenced by a statistically significant increase in fluorescence intensity compared with Vehicle control, and H105A decreases CSE-induced cell death at both tested concentrations. CSE+H105A(1nM) = 848.9 ± 51.0 and CSE+H105A(20nM) = 919.2 ± 46.0, vs. CSE alone = 1447.7 ± 175.3 and CSE+R99A(20nM) = 1283.2 ± 67.6. Mean ± SEM; **** p<0.0001, One-way ANOVA. (E) Representative photomicrographs of retinal organoid sections stained with TUNEL to assess apoptosis-induced DNA fragmentation. Dashed lines demarcate ONL. Scale Bar: 50 μm. (F) Histogram of quantification of TUNEL staining shows a statistically significant increase in cell death in CSE-treated ROs compared to vehicle controls, and a statistically significant decrease in CSE-induced cell death by H105A treatments. Relative TUNEL(+) area: CSE+H105A(1nM) = 2.77 ± 0.25 and CSE+H105A(20nM) = 2.35 ± 0.23, vs. CSE alone = 7.36 ± 0.25 and CSE+R99A(20nM) = 5.27 ± 0.50. Mean ± SEM; **** p<0.0001, One-way ANOVA.

Using 3D-ARQ, a technique we previously developed to quantify fluorescence intensity in live 3D ROs (*44*), we evaluated PSVue®-794 staining in peptide-treated ROs and found that H105A treatments significantly decreased PS externalization in CSE-exposed ROs, relative to CSE alone and CSE+R99A treatments (**Fig. 8C**). Moreover, H105A prevented CSE-induced cell-death in human ROs as shown using ethidium homodimer, a membrane-impermeable fluorescent dye that binds to DNA in dead or dying cells. H105A treatment significantly decreased CSE-induced cell death relative to treatments with CSE alone and CSE+R99A (**Fig. 8D**). Consistently, to detect DNA fragmentation occurring in late apoptosis, TUNEL assays were performed in cryosections of D180 ROs treated as described above. H105A treatments significantly decreased the number of TUNEL-positive nuclei in the ONL of ROs undergoing CSE-induced cell death to levels as observed for the untreated ROs (Veh) (**Figs. 8E, 8F**).

Altogether, our findings demonstrated that the H105A peptide promoted the survival of human retinal photoreceptors in ROs, highlighting again its translational potential for medical and therapeutic applications related to retinal diseases.

## DISCUSSION

Building upon our previous findings on structure-activity relationships of PEDF, in this study we investigated the potential use of small peptides derived from the neurotrophic active region of PEDF delivered as eye drops as a treatment option for RP. We report that 17-mer and H105A peptides, each composed of 17 amino acid residues, act as neurotrophic agents that preserve photoreceptor viability, morphology, and function in models of retinal degeneration mimicking human RP. Our study demonstrates that these peptides, when locally delivered to the eye, achieve sufficient retinal bioavailability to effectively target PEDF-R, the PEDF receptor crucial for photoreceptor survival (*27,28*). Consequently, they block cell death pathways associated with retinal diseases. Key findings supporting these conclusions include: 1) the 17-mer and H105A peptides target murine photoreceptors and activate PEDF-R when delivered via eye drops or intravitreal AAV2-viral injections; 2) these peptides block PS externalization and DNA fragmentation and increase the BCL2/BAX ratio in photoreceptors of mutant mice undergoing spontaneous cell death; and 3) the peptides improve both the morphology and function of degenerating murine photoreceptors. Peptides showing poor penetration (e.g., 29-mer, **Fig. S4**) and those lacking PEDF-R affinity (e.g., R99A) were ineffective. Additionally, the results obtained using a human retinal organoid model undergoing oxidative stress-induced cell death provide proof-of-concept for the efficacy of the H105A peptide in preventing damage to human retinal cells, further supporting its translational potential for retinal degenerative diseases. Altogether, the findings strongly emphasize the potential of peptide therapeutic approaches for preserving or enhancing vision in retinal degenerative diseases.

The significance and potential impact of our study are grounded on its translational promise as it demonstrates the ability of 17-mer and H105A peptides to stabilize the progression of photoreceptor degeneration in preclinical murine models of human retinal diseases. These peptides show potential to overcome the challenging genetic heterogeneity characterizing RP as demonstrated in distinctive models: the *rd10* containing an alteration in the visual transduction enzyme Pde6, and the *Rho^P23H/+^* characterized by an alteration in the light receptor rhodopsin protein, modelling different types of human RP; and the *rd10/Serpinf1s^−/^*, a more aggressive variant of the *rd10* model lacking PEDF; as well as human hiPSCs-derived retinal organoids undergoing oxidative stress, a feature of RP (*12, 16, 18*). Not only did the peptides act on preserving photoreceptor morphology but also on improving the light responses of both types of photoreceptors, rods, and cones. These genetically distinct models have common and unique retinopathies, suggesting that the H105A peptide appears to have broad applicability for a range of human retinal diseases involving photoreceptor degeneration. This is primarily due to its ability to target intracellular calcium, a common feature in photoreceptor degeneration (*18*). We envision that the H105A peptide would be a beneficial therapeutic agent for other diseases that result in photoreceptor degeneration, including but not limited to other RP types, age-related macular degeneration, choroideremia, Usher syndrome, Stargardt disease, diabetic retinopathy, etc. Further assessments of H105A peptides will be required to identify potential benefits for these and other diseases characterized by photoreceptor degeneration.

The novelty of this study is broadened by the neurotrophic H105A peptide therapeutic delivered via eye drops, which offers a high safety profile and multiple delivery options. Peptide eye drops are an attractive alternative to gene-therapy approaches administered via intraocular injections. Eye drop administration is simpler, less invasive, and less likely to damage the retina, potentially improving patient compliance. Although several attempts to develop eye drop delivery for retinal disorders have been unsuccessful due to membrane barriers and drug digestion at the eye surface (*5*), there are promising precedents. For example, a small cyclic peptide of 5 residues, Vasotide, delivered via eye drops effectively reached the posterior ocular tissues to inhibit retinal and choroidal angiogenesis in mice and monkeys (*45*). Additionally, in a pilot study, topical administration of nerve growth factor eye-drops in RP patients was found to be safe with some patients experiencing improvements in visual performance (*46*). Our findings show successful delivery of therapeutic peptides via eye drops, with transfer of 17-mer and H105A peptides into the posterior retina, where the RP photoreceptors undergo degeneration. We speculate that eye drop delivery could be enhanced using carrier systems like nanoparticles, or liposomes, which could improve stability and bioavailability of PEDF neurotrophic peptides. Moreover, peptide eye drops could serve as a preemptive treatment while waiting the development of specific gene-targeted therapies for individual patients. Other peptides, derived from the PEDF antiangiogenic region (34-mer), which is distinct from the neurotrophic region, have successfully decreased experimental choroidal and retinal neovascularization in mice and rabbits when injected IVT as free peptides, mixed with collagen or encapsulated in nanoparticles (*23, 47–49*). The second novelty lies in the gene therapy approach delivered by IVT injection. AAV2 IVT-based gene therapy is already in clinical trials for retinal ganglion cell diseases. Our approach, however, targets photoreceptors through an engineered cDNA construct that allows the therapeutic agent to be secreted in the retinal environment. This method avoids the risk of retinal detachment and provides broad retinal coverage, unlike localized subretinal injections. While PEDF peptides 17-mer and H105A show promise in preclinical studies, further research is needed to optimize delivery methods and establish their long-term safety and efficacy. Nevertheless, these observations represent a significant advancement in understanding how structure-activity relationships of a naturally occurring bioactive protein can facilitate therapeutic drug development.

In summary, we present novel compounds and methods for treating retinal degeneration using PEDF-derived peptides. This modality of peptide therapeutics is promising for photoreceptor death prevention in several retinopathies, and topical ocular peptide delivery has potential for translation into safer, less-invasive applications to stabilize and delay the progression of pathological neurodegeneration in retinal disorders in a large proportion of RP patients.

## METHODS

### Animals

Experimental animals *rd10* mice (from Jackson Laboratories) and *rd10/Serpinf1^−/−^* (*16*), (*rd10* mice crossed with *Serpinf1 null* mice, double mutant mice) of P0-P25 days old were maintained in the animal facility of the National Institutes of Health, USA. Experimental animals *Rho^P23H/+^* (*17*) from P0 to P180 were maintained in the animal facility CSSI (Centro Servizi Stabulario Interdipartimentale, Italy). All animals were on normal chow diet and a 12-h light/dark cycle. All the experimental procedures were approved by the National Eye Institute Animal Care and Use Committee and the Ethical Committee of University of Modena and Reggio Emilia and by the Italian Ministero della Salute (150/2021-PR) and were performed as per guidelines of the Association for Research in Vision and Ophthalmology statement for the Use of Animals in Ophthalmic and Vision Research and in accordance with the ARRIVE guidelines.

We did not select either males or females in our studies because the murine models here used have not been reported to show differences in the progression of the disease between sexes, as also reported in patients with mutations in the PDE6 enzyme or P23H mutation in rhodopsin. Therefore, based on the “reduction” 3R requirement for animal studies, we did not exclude animals from the study, and all the bred animals participated in the study.

### Peptides and proteins

The peptides were chemically synthesized and purified to ≥95% purity (Biosynthesis Inc. and LifeTein). They were conjugated to Alexa Fluor 488 dye to produce fluorescently labeled peptides (BioSynthesis, and LifeTein). The peptide sequences used in the study are shown in **Table S1**.

Lyophilized peptides were dissolved in sterile HBSS at 1 mg/ml and stored at −20 °C. Before the peptides were used, the peptide solution was brought to room temperature and subjected to centrifugation (Eppendorf, Centrifuge 5430R) at 20,800 g for 10 min at 4°C to clear and separate particulate material. In case a precipitate was found, the pH of the solution was adjusted to neutral pH with either NaOH or HCl and all peptides were soluble. The concentration of peptide solutions was determined using spectrophotometry (NanoDrop, Thermo Fisher Scientific). The stability of peptide H105A against temperature was performed and summarized in **Table S2**.

To obtain the PEDF-R protein, the following procedure was followed. For plasmid construction, the plasmid used for expression was pE-SUMOstar-PEDF-R[1-288] C-terminal TwinStrep tagged. This plasmid contained the coding sequence for the human PEDF-R ORF between positions 1 and 288. For transformation: The plasmid pE-SUMOstar-PEDF-R[1-288] and pGro7(groES-groEL) were co-transformed into *Escherichia coli* Rosetta(DE3) to obtain colonies for recombinant protein expression, as described before (*50, 51*). For cell harvesting: Cells were grown to the desired density and then harvested by centrifugation. For cell lysis: The harvested cells were resuspended in buffer A (50 mM Tris-HCl, pH 8.0, 500 mM NaCl, 20% Glycerol, 1 mM TCEP, 0.1% IGEPAL CA-630, 0.5% (W/V) CHAPS and 10 mM Imidazole) containing EDTA-free protease inhibitor cocktail. The cells were then lysed, and the lysates were cleared by centrifugation (33,745 g, 30 min, 4°C). For affinity chromatography: The cleared lysate supernatant was subjected to two-step affinity chromatography using a HisTrap HP column followed by a StrepTrap XT column. This process was performed using an automated ÄKTA pure™ chromatography system (Cytiva). Fractions containing the recombinant protein were collected and pooled. The pooled fractions were concentrated, resulting in a recombinant PEDF-R[1-288] protein that was >90% pure, soluble, and active. This protocol ensured the successful purification of the recombinant PEDF-R protein, which can be used for further experiments and analyses.

### Retinal extracts

Protein extracts were obtained from dissected mouse retinas in RIPA Lysis and Extraction Buffer (Thermo Fisher Scientific, cat. 89900) containing protease inhibitors (Pierce Protease Inhibitor Tablets, Thermo Fisher Scientific, cat. A32963) at 80 µl per retina, as described previously (*28*).

### Eye drop administration

In a sterile hood, aliquots of the peptide solutions were prepared from the stocks. Administration to each mouse eye was with a volume of 5 µl at the indicated concentrations and regimen of treatment (see schemes in figures). The eye drops were administered by the same investigator throughout the study for consistency.

### Quantification of peptides in the retina

Each AlexaFl488-labeled peptide solution was administered via 5 µl of eye drops containing peptide at 1 mg/ml concentration per eye of wild type mice. At end point, retinal protein extracts were prepared from each enucleated eye. A total of 30 µl of each freshly prepared soluble retinal extract was added to a well in 96-well plate in triplicates. Fluorescence was determined using a fluorometer (POLARstar OPTIMA) using a wavelength at 485 nm for excitation and at 520 nm for emission. Retinal extract without peptide was used as a background control set at zero. The amount of peptide in the retina was determined from the fluorescence and using standard curves (**Fig. S5A**).

#### Standard curves of Alexa Flour-488 labeled peptides

Retinal extracts from three C57Bl/6J mouse eyes were prepared (see Methods), pooled and used to dilute the labeled peptides. Solutions of Alexa Flour-488 labeled-17-mer, -H105A and -R99A peptides at concentrations that ranged between 0.0 µg/ml and 0.0625 µg/ml were prepared in retinal extracts. A total volume of 30 µl of each was added to a well of a 96-well plate. The fluorescence in the retinal extract was determined with a plate reader of a fluorometer (POLARstar OPTIMA) using a wavelength at 485 nm for excitation and at 520 nm for emission. Plots were prepared from the fluorescence as a function of peptide concentration using GraphPad. Standard curves were used to quantify the labeled peptides 17-mer, H105A and R99A in the retina extracts (see **Fig. S5A**). AlexaFl488 labelled peptides 17-mer and H105A exhibited neuroprotective activity, as assessed by ERG, indicating they were functional peptides, but R99A was not (**Fig. S5B**).

### Phospholipase A_2_ activity assay

Phospholipase A_2_ (PLA_2_) activity was determined using EnzChek™ Phospholipase A2 Assay Kit (Thermo Fisher Scientific, cat. E10217) according to the manufacturer’s instructions and as described before (*28*). Protein samples (retinal extracts or PEDF-R[1-288], as described above) were mixed with peptides and preincubated at room temperature for 30 min and diluted up to 50 µl before enzymatic evaluation. Briefly, 50 µl of the sample were transferred to a microplate well (96-well plate). A lipid mixture was prepared by mixing 15 µl 10 mM dioleoylphosphatidylcholine (DOPC), 15 µl 10 mM dioleoylphosphatidylglycerol (DOPG) and 15 µl 1 mM PLA_2_ substrate. Then, a substrate-liposome was prepared by mixing 25 µl lipid mixture with 2.5 ml 1X PLA_2_ reaction buffer by stirring. A total of 50 µl of substrate-liposome mixture was added to each microplate well containing the sample, and incubated at room temperature for 10 minutes, protected from light. The final reaction volume was 100 µl. Fluorescence emission was measured at 515 nm with excitation at 460 nm. The final concentration of PEDF-R[1-288] was 60 nM in the reaction mixture.

### *In vivo* photoreceptor degeneration assessment in *rd10* and *rd10 x Serpinf1^−/−^* murine models

Fluorescence fundoscopy with a cell death probe was chosen as a new and simple method for monitoring photoreceptor cell death in alive animals (*35*). A fluorescent probe, bis (zinc^2+^-dipicolylamine)-550, (PSVue^®^-550) (Molecular Targeting Technologies Inc, cat. P-1005) was reconstituted in Hanks’ Balanced Salt Solution (HBSS, Quality Biological, cat. 114-062-101) according to the manufacturer’s instructions and as previously described (*28, 35*). This probe is suitable for *in vivo* use to specifically and transiently label dying photoreceptors, such as in living rats with retinal degeneration the Royal College of Surgeons rat model, in a non-invasive and non-toxic method without the need of intraocular injection (*35*). Briefly, a stock of 1 mM solution of PSVue^®^-550 was stored in the dark at 4°C and used within 14 days directly as eye drop. After 3-4 h of light onset each mouse received eye drops of 5-10 μl of 1 mM PSVue^®^-550 in the left eye and HBSS in the right eye as control. The eye drop volume was chosen such that the eye cavity was filled without spillage outside the eye, and it was modified depending on eye size. Eyes of mice that did not receive the probe were used a background controls. To obtain *in vivo* retinal images of fundi, at 24 h after the probe was administered, the mice were anesthetized, and pupils were dilated with 1% tropicamide for 5 min and kept hydrated with GenTeal. Fluorescence was imaged using a retinal imaging microscope (Micron III, Phoenix Research Labs) using an FF02-475/50 nm excitation filter (Semrock, Inc.).

To determine the natural history of photoreceptor death of *rd10* and *rd10/Serpinf1^−/−^* mice, we assessed photoreceptor cell death in live mice using the fluorescent probe termed PSVue® 550 to monitor PS externalization by fluorescence fundoscopy (see **Fig. S1**). Given the reports about the retinas of *rd10* mice starting to show histological changes at P16, and with a peak in the number of TUNEL+ nuclei in the ONL at P17-18 (*13*), we monitored PS externalization in photoreceptors of *these* mice undergoing degeneration before death onset (**Fig. S1A**). Eye drops of 1 mM PSVue® 550 solution in HBSS were administered to left eyes and HBSS (vehicle) to their contralateral right eyes of each mouse at ages P16, P20, P22 or P24. PS externalization was determined by fluorescence fundoscopy 24-h post-administration.

### Retinal cross sections, histology, and immunofluorescence

Both eyes were oriented before enucleation to allow sectioning along the dorsal/ventral axis. Enucleated eyes from *rd10* and *rd10 x Serpinf1^−/−^* mice were incised in the cornea and fixed in 2.5% glutaraldehyde for 20 min and then in 10% neutral buffered formalin at 4°C for at least 48 h, followed by paraffin embedding, sectioning, and staining with either H&E or immunofluorescence procedures, as previously described (*28*). For fluorescence microscopy, antibodies, and 0.1 µg/ml DAPI were used to immunostain and nuclei counterstain the sections, respectively. Antibodies used in the study are in **Table S3**. Images were acquired using a ZEIS 700 confocal microscope. Fluorescence intensity was quantified using ImageJ as previously described (*14, 26*).

Eyes from *Rho^P23H/+^* mice eyes were enucleated and fixed in Davidson’s fix (Formaldehyde 8% (Merck, cat. F8775); Ethanol 32% (Carlo Erba, cat. 414608); Acetic Acid 10% (Merck, cat. 695092) over-night. After embedding in paraffin, 14 μm sections were collected. For TUNEL, sections were treated with 0.1 M Citrate Buffer pH 6 (Tri-sodium Citrate 10 mM (Carlo Erba, cat. 368007); Citric Acid 0.2 mM (Merck, cat: 251275) in microwave for 5 minutes, then labeling was performed for one hour at 37°C using a In Situ Cell Death Detection Kit (Merck, cat. 12156792910). Rod photoreceptors were labeled with anti-RHO 1D4 antibody followed by Alexa Fluor 568 anti-mouse secondary antibody (see **Table S3**). Cone photoreceptors were labeled with peanut agglutinin (PNA, Vector Laboratories, cat. FL-1071). GFP localization in control transduced retinas was identified with specific anti-GFP antibodies and Alexa Fluor 488 anti-mouse secondary antibody (see **Table S3**). For nuclear staining, 0.1 μg/mL DAPI was used. Slides were mounted with Mowiol 4–88 (Merck) and images acquired with a Zeiss Axio Imager A2 fluorescence microscope. To count TUNEL positive cells the RETINA Analysis Tool kit for ImageJ was used (*52*).

To generate spider plots, the ONL thickness was measured from images acquired with light microscopy of the retina sections stained with H&E or images of retina sections stained with DAPI with a dorsal-ventral orientation. The ONL thickness in the entire retina sections was measured at 5 different positions from both sides of the optic nerve (ON) using ImageJ, as described before (*28*). Alternatively, for DAPI stained images, a customed macro implemented in ImageJ was used to select only the ONL.

All images from different mice of each genotype and age were collected with identical magnification, gain and exposure settings, and the same retinal region. Each experimental group included at least 5 eyes.

### Retinal functional assessment of *rd10* and *rd10/Serpinf1^−/−^* mice

Electroretinogram (ERG) for *rd10* and *rd10/Serpinf1^−/−^*mice were recorded in both eyes of each animal using an Espion E2 system with ColorDome (Diagnosys LLC) with a heated surface and mice were dark-adapted overnight as previously described (*29*). Mice were placed on the heated surface and electrodes were placed in the mouth and a subdermal platinum needle electrode was placed in the back of the mouse to serve as ground. Responses were elicited with increasing light impulses with intensity from 0.0001 to 10 cd-seconds per meter squared (cd.s/m^2^). Amplitudes for a-wave were measured from stimulus to the trough of the a-wave and b-wave amplitudes were measured from a-wave to b-wave trough or peak. The values from the a and b waves for both eyes were exported to Microsoft Excel to obtained amplitude values for each mouse.

### Viral delivery and evaluation of photoreceptor degeneration in the *Rho^P23H/+^* mouse model

The cDNA encoding H105A was cloned in frame with the signal peptide from Interferon beta (*53*), to allow secretion of the H105 peptide, in a pAAV2 vector with a CMV promoter resulting in AAV-H105A. The AAV-H105A and AAV-GFP, as control, viruses were generated by InnovaVector (*54*). For AAV2 delivery, *Rho^P23H/+^* pups at P5 were anesthetized for 2 minutes in ice. After eyelid opening, 0.5 µl of AAV-H105A (1.9 × 10^12^ genome copies (GC)/ml) (one eye) or AAV-GFP (in the contralateral eye, 4.5 × 10^12^ GC/ml) were intravitreally injected with a 34 GA needle Hamilton syringe. A topical cortisone and antibiotic unguent (TobraDex, Alcon, 0.3% tobramycin + 0.1% dexamethasone) was applied immediately after injection and pups were allowed to recover on a warm pad before reintroduction in the cage.

### AAV-17-mer[H105A] mRNA expression

Retinas were microdissected and lysed, and total RNA was purified with RNeasy Mini Kit (Qiagen, Hilden, Germany, catalogue number 74104). 200 ng of total RNA were retrotranscribed with iScript cDNA Synthesis Kit (Biorad, Hercules, CA, USA, catalogue number 1725035) following manufacturer’s instruction, and cDNA was used for PCR using the following primers: H105A forward (TGCAGAAGTTGGTCGTGAGG), and H105A reverse (GGATTCTGTTCGCTGGATCC). The amplicons were resolved by electrophoresis in a 2% agarose gel stained with ethidium bromide (0.5 ug/ml, Merck), and MW markers 100 bp DNA ladder (Promega, G2101).

### AAV-17-mer[H105A] protein expression

To detect the transduced H105A peptide, we used a custom-made monoclonal recombinant Fab antibody raised against H105A (Bio-Rad AbDSerotec GmbH, Puchheim, Germany). The custom-made antibodies were produced by HuCAL® technology with a FLAG-tag for detection. To assess antibody labelling of H105A, COS-7 cells were infected with AAV-H105A at 1.9 GC/ml. At 48 hours post-transduction, cells were fixed in 2% PFA for 10 minutes and, after blocking with 3% BSA in PBS, were incubated with 10 mg/ml anti-H105A antibody (clone number AbD57564pao) diluted in 1% BSA and 0.1% Tween 20 at 4°C for 16 hours, followed by incubation with secondary antibody, anti-FLAG antibody (Merck, catalogue number F3165) diluted 1:100 in 1% BSA and 0.1% Tween 20 at 25°C for 2 hours and Alexa Fluor 568 anti-mouse as tertiary antibody (Thermo Fisher Scientific, catalogue number A-11004) diluted 1:1000 in 0.1% BSA at 25°C for one hour. The specificity of the custom antibody to the H105A peptide was assessed by pre-incubation with 400 µM H105A peptide at 25°C for 1 hour before using it for immunodetection. Production of the recombinant H105A peptide in retinal cells after AAV-H105A IVT was evaluated by immunofluorescence with the same protocol. Slides were mounted using Mowiol 4-88 (Merck) and images were acquired using an SP8 confocal microscope (Leica, Heidelberg, Germany) with a 40X oil objective, equipped with white-light laser.

Viral expression assessments are shown in **Figure S3**.

### Retinal functional assessment in *Rho^P23H/+^* mice

Electroretinograms for *Rho^P23H/+^* mice were recorded in both eyes of each animal. Six-month-old mice were anesthetized and dark-adapted overnight as previously described (*55*). Mice were placed inside a 30 cm diameter Ganzfeld sphere, and coiled gold electrodes were placed in the corneas moistened by a thin layer of gel (Lacrinom, Farmigea, Italy) and the reference electrode (earth) was placed in the scalp. Six calibrated neutral density filters were utilized to regulate the intensity of the light stimulation, which was performed using a white light electric flash (SUNPACK B3600 DX, Tecad Company, Tokyo, Japan). Scotopic ERG responses were registered with increasing light stimuli (0.0041 to 377.2 cd.s/m^2^, using 0.6 log units increments) with an inter-stimulus interval spanning from 20 s for dim flashes to 45 s for the brightest flashes. After 15 minutes from background onset, isolated cone (photopic) components were registered by superimposing the test flashes (0.016 to 377.23 cd.s/m^2^) over a stable background of saturating intensity for rods (30 cd/m^2^). From zero to the b-wave peak, the amplitude of the scotopic and photopic b-waves was measured.

### Stem cell-derived retinal organoid generation

We utilized the human-derived induced pluripotent stem cell (hiPSC) cell line A18945 (Gibco), which was sourced from female cord blood. This hiPSC line was cultured on an extracellular Matrigel basement membrane matrix (Corning, 354230) and maintained in mTeSR1 media (StemCell Technologies) until they were 60-70% confluent and ready for differentiation.

Human iPSCs were differentiated into retinal organoids (ROs) following the protocol by Zhong et.al (*39*). Briefly, on Day 0, hiPSC colonies were dissociated with dispase and grown in a suspension to form embryoid bodies, and slowly transitioned to neural induction media (DMEM/F12 (1:1), 1% N2 supplement (Invitrogen), 1 x minimum essential media-nonessential amino acids (NEAAs), 2 µg/ml heparin (Sigma)). On Day 7, EBs were seeded on Matrigel coated plates and maintained in neural induction media. On Day 16, the media was switched to the retinal differentiation medium (DMEM/F12 (3:1), 2% B27 (without vitamin A, Invitrogen), 1x NEAA and 1% antibiotic–antimycotic (Gibco)). Retinal domains were identified and lifted manually with tungsten needles between Day 21 and Day 26 under a phase contrast microscope. On day 30, the culture medium was transitioned to DMEM/F12 (3:1) with 2% B27, 1x NEAA, 1% antibiotic–antimycotic, 10% fetal bovine serum (FBS; Gibco), 100 mM Taurine (Sigma), and 2 mM GlutaMAX (Invitrogen). Additionally, 1 µM retinoic acid supplement was introduced daily starting from day 63. On day 91, the medium was modified to DMEM/F12 (1:1) supplemented with 1% N2, 1 x NEAAs, 1% antibiotic–antimycotic, 10%FBS, 100 mM Taurine and 2 mM GlutaMAX, and the concentration of retinoic acid supplementation was reduced to 0.5 µM for the remainder of the culture period. The use of hiPSCs in this study conforms to the University of Colorado Institutional Biosafety Committee standards.

### RO culture treatments

On Day 180, live ROs were individually plated into transparent U-bottom 96-well plates, washed three times with PBS (Gibco, 14190-144) and treated according to each of five experimental conditions: Vehicle Control (DMSO), Cigarette Smoke Extract (CSE) (500 µg/ml) (Murty Pharmaceuticals, NC1560725), CSE with R99A (20 nM), and CSE with H105A (1 nM and 20 nM), for 24 hours at 37°C in a CO_2_ incubator.

### Quantitative assessment of cell death in live ROs

After treatment, ROs were washed in PBS (Gibco, 14190-144) and the auto-fluorescent background was measured by 3D-automated reporter quantification (3D-ARQ) as previously described (*44*), using a Tecan Spark Plate reader (SparkControl v2.3). The Z-position was determined manually by selecting the individual well and fluorescent intensity was measured using the optimized parameters below. ROs were then incubated with 10 µM PSVue-794 (Molecular Target Technologies, P-1001) for 1 hr at 37°C, or with 4 µM of Ethidium Homodimer (Thermo Fischer Scientific, L3224) for 45 minutes at 37°C. After incubation, ROs were washed three times with 1X PBS, and fluorescence intensity was measured by 3D-ARQ using the following parameters: Mode: Fluorescent top reading; Wavelength for PSVue-794: Excitation: Emission 730:820 nm, Excitation bandwidth: 10 nm, Emission bandwidth: 15nm; Wavelength for Ethidium Homodimer: Excitation: Emission 530:630 nm), Excitation bandwidth: 5 nm, Emission bandwidth: 5nm; Gain: 160, Manual; Number of flashes: 20; integration time: 40 µs; lag time 0 µs. Clear DMEM media was used as a blank. Results were analysed after background subtraction; n=19-24 ROs per condition.

### TUNEL Staining

D180 ROs were treated with each of the 5 experimental conditions, as described above (n=5 for each condition), washed with PBS three times, and fixed with 4% paraformaldehyde for 10 minutes at room temperature. ROs were then subjected to overnight incubation in 6.75%, 12.5% and 25% sucrose solutions in PBS, embedded in Optimal cutting temperature compound (OCT compound, Fisher Healthcare #4585): Sucrose (1:1) solution, and stored at −80°C. Cryosections of 10 μm were prepared using a Leica Cryostat. Slides were dried and washed with PBS for 15 minutes. Subsequently, slides were rinsed three times with deionized water for 5 minutes and incubated in 1x Terminal Transferase (TdT) buffer containing 1X buffer 4 (NEB) and 2.5 μM CoCL_2_ for 10 minutes at room temperature in a humid chamber. For labelling, TdT buffer, biotinylated 16 -dUTP and TdT were added to each slide and incubated for 1 hour at 37°C in a humid chamber. The reaction was stopped by incubating slides in 2XSSC (Research Products International Corp. #S24022-1000.0) for 15 minutes at room temperature followed by a wash with PBS. Blocking was performed using 2% BSA in PBS for 10 minutes at room temperature, and slides were incubated with streptavidin CY3 (ThermoFisher Scientific #438315) for 45 minutes at room temperature in a humid chamber. Subsequently, slides were washed 3 times with PBS, counterstained with DAPI for 10 minutes, washed with PBS and DI water and mounted using FluoroMount solution (SouthernBiotech #0100-01). Fluorescence images were acquired with a Nikon C2 Confocal Microscope A1 (Minato City, Tokyo, Japan) and stitched together using Nikon Elements Software. Images were analysed with Fiji. The relative area of positive immunostaining was calculated as a percentage of total retinal area.

### Statistical analyses

We performed each *in vivo* experiment at least three times and used representative data per treatment and age in the calculations.

Data were analyzed using GraphPad Prism version 9.5.0 for Windows (GraphPad Software). All experimental groups for *rd10* and *rd10/Serpinf1^−/−^* mice were compared to each other using a two-tailed unpaired Student t test or one-way ANOVA using Dunette’s multiple comparison as we compared the mean of the control group with the other groups. All groups are shown as mean ± SD. *p*-values lower than 0.05 were considered statistically significant.

For Spider plot analysis for *Rho^P23H/+^* mice, all experimental groups were compared to each other using Two Way ANOVA with Šídák’s multiple comparisons test as we compared the mean of the control group with the other groups. For ERG analysis One-way ANOVA was used. All groups are shown as mean ± SEM. *p*-values lower than 0.05 were considered statistically significant.

Statistical analyses for human ROs were performed with GraphPad Prism 10.1.0 Software using one-way ANOVA for multiple comparisons. *p*-values less than 0.05 were considered statistically significant. Bar graphs depict the means ± SEM with individual values plotted.

## Acknowledgments

We thank Megan Kopera for assistance with animal husbandry, Haohua Qian for assistance and usage of the Visual Function Core, Maria Campos for service of the Histology Core and Robert Fariss for access to the NEI bioimaging Core. We thank K. Palczewski for the *Rho^P23H/+^*mouse model. We thank K. Bharti for proofreading this manuscript.

## Funding

The Intramural Research Program of the National Eye Institute, National Institutes of Health, United States of America. Project #EY000306 (SPB)

The Prevention of Blindness Society (SPB)

Fondazione Telethon. Project #GGP19113 (VM)

National Center for Gene Therapy and Drugs based on RNA Technology” cod. Progetto CN00000041 and “Health Extended ALliance for Innovative Therapies, Advanced Lab-research, and Integrated Approaches of Precision Medicine - HEAL ITALIA” tematica 6 “Innovative diagnostics and therapies in precision medicine” cod. Progetto PE0000019 PIANO NAZIONALE DI RIPRESA E RESILIENZA (PNRR) – MISSIONE 4 “Istruzione Ricerca” COMPONENTE 2, “Dalla ricerca all’impresa” INVESTIMENTO 1.4, “Potenziamento strutture di ricerca e creazione di “campioni nazionali di R&S” su alcune Key enabling technologies”, finanziato dall’Unione europea – NextGenerationEU (VM and AB)

The *CellSight* Development Fund (NV)

A Challenge Grant to the Department of Ophthalmology at the University of Colorado from Research to Prevent Blindness (NV)

## Author contributions

A.B.C contributed Methodology, Investigation, Formal analysis, Visualization, Writing - original draft, Writing - review & editing.

A.B. contributed Investigation, Formal analysis, Visualization, Funding acquisition, Writing - review & editing.

S.P. contributed Investigation, Formal analysis, Visualization, Writing - review & editing.

S.D. contributed Investigation, Visualization, Writing - review & editing.

I.P contributed Formal analysis, Writing - review & editing.

E.A. contributed Investigation.

F.C. contributed Investigation.

C.G. contributed Formal analysis, Writing - review & editing.

N.V. contributed Conceptualization, Methodology, Visualization, Funding acquisition, Supervision, Writing - review & editing.

V.M. contributed Conceptualization, Methodology, Funding acquisition, Supervision, Writing - review & editing.

S.P.B. contributed Conceptualization, Methodology, Visualization, Funding acquisition, Project administration, Supervision, Writing - review & editing.

## Competing interests

US Government and University of Modena and Emilio Reggio, Italy. S. P. Becerra, A. Bernardo-Colón, Valeria Marigo, A Bighinati. US Application No. 63 430 251 NIH Ref No E-028-2023-0-US-01 and international application PCT/US2023/064947, filed on March 24, 2023. The specific aspect of the manuscript covered in the patent applications is on PIGMENT EPITHELIUM-DERIVED FACTOR PEPTIDES AND USE FOR TREATING RETINAL DEGENERATION.

## Data and materials availability

All data are available in the main text or the supplementary materials.

## List of Supplementary Materials

Figures S1 to S5

Tables S1 to S3

## Notes

### Competing Interest Statement

The authors have declared no competing interest.

