## Supplementary Materials for "H105A peptide eye drops promote photoreceptor survival in murine and human models of retinal degeneration"

### Figure S1.

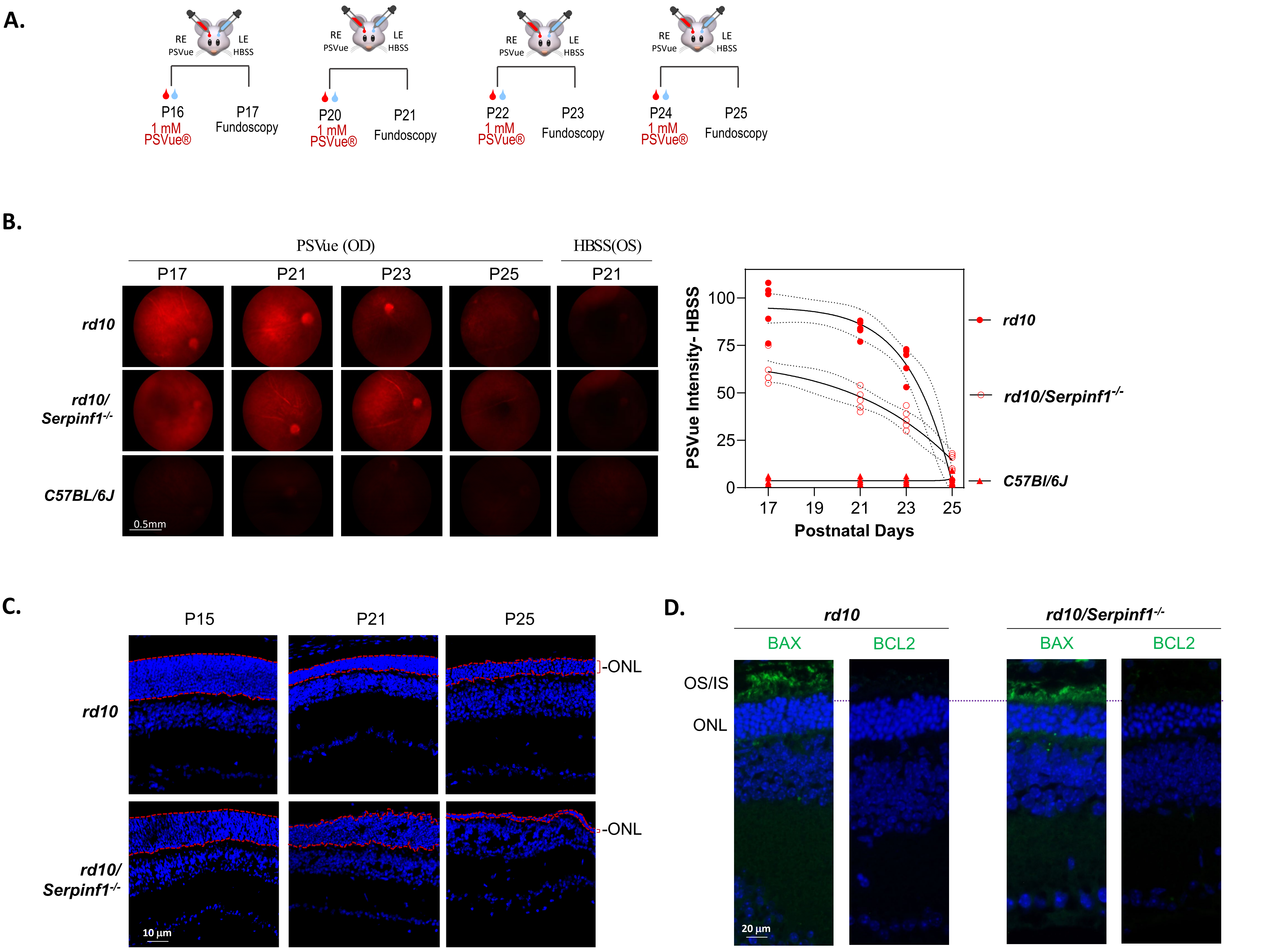
A.
RE PSVue
LE HBSS
P16
1 mM PSVue®
P17 Fundoscopy
P20
1 mM PSVue®
P21 Fundoscopy
P22
1 mM PSVue®
P23 Fundoscopy
P24
1 mM PSVue®
P25 Fundoscopy
B.
PSVue (OD)
HBSS(OS)
P17
P21
P23
P25
P21
rd10
rd10/ Serpinf1-/-
C57BL/6J
0.5mm
PSVue Intensity- HBSS
100
75
50
25
0
17
19
21
23
25
Postnatal Days
rd10
rd10/Serpinf1-/-
C57Bl/6J
C.
P15
P21
P25
rd10
rd10/ Serpinf1-/-
-ONL
-ONL
10 µm
D.
rd10
rd10/Serpinf1-/-
BAX
BCL2
BAX
BCL2
OS/IS
ONL
20 µm

**Figure S2.**

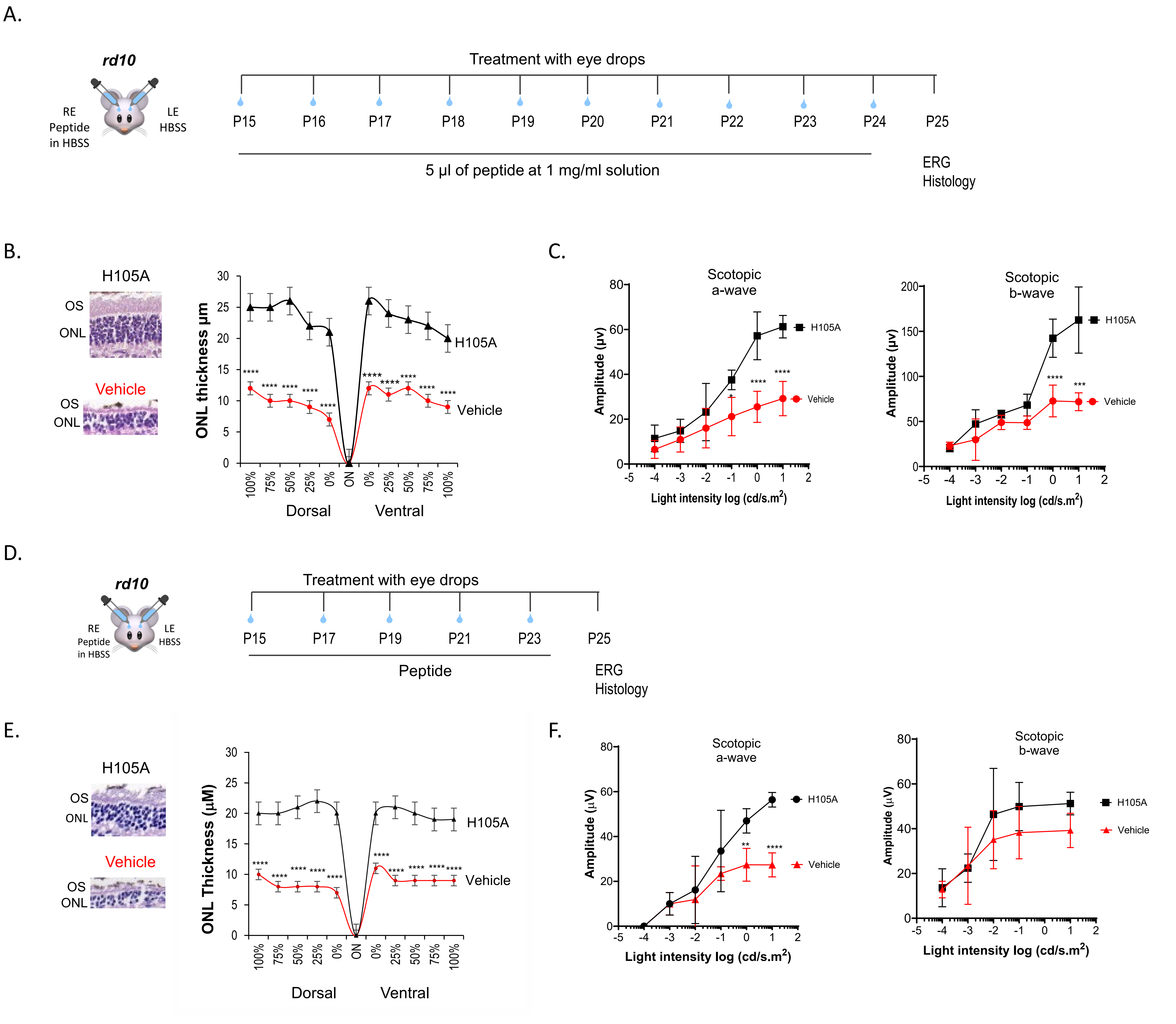
A.
rd10
RE Peptide in HBSS
LE HBSS
Treatment with eye drops
P15
P16
P17
P18
P19
P20
P21
P22
P23
P24
P25
5 µl of peptide at 1 mg/ml solution
ERG
Histology
B.
H105A
OS
ONL
Vehicle
OS
ONL
ONL thickness µm
H105A
Vehicle
100%
75%
50%
25%
0%
ON
0%
25%
50%
75%
100%
Dorsal
Ventral
C.
Scotopic a-wave
Amplitude (µv)
H105A
Vehicle
Light intensity log (cd/s.m2)
Scotopic b-wave
Amplitude (µv)
H105A
Vehicle
Light intensity log (cd/s.m2)
D.
rd10
RE Peptide in HBSS
LE HBSS
Treatment with eye drops
P15
P17
P19
P21
P23
P25
Peptide
ERG
Histology
E.
H105A
OS
ONL
Vehicle
OS
ONL
ONL Thickness (µM)
H105A
Vehicle
Dorsal
Ventral
F.
Scotopic a-wave
Amplitude (µV)
H105A
Vehicle
Light intensity log (cd/s.m2)
Scotopic b-wave
Amplitude (µV)
H105A
Vehicle
Light intensity log (cd/s.m2)

**Figure S3.**

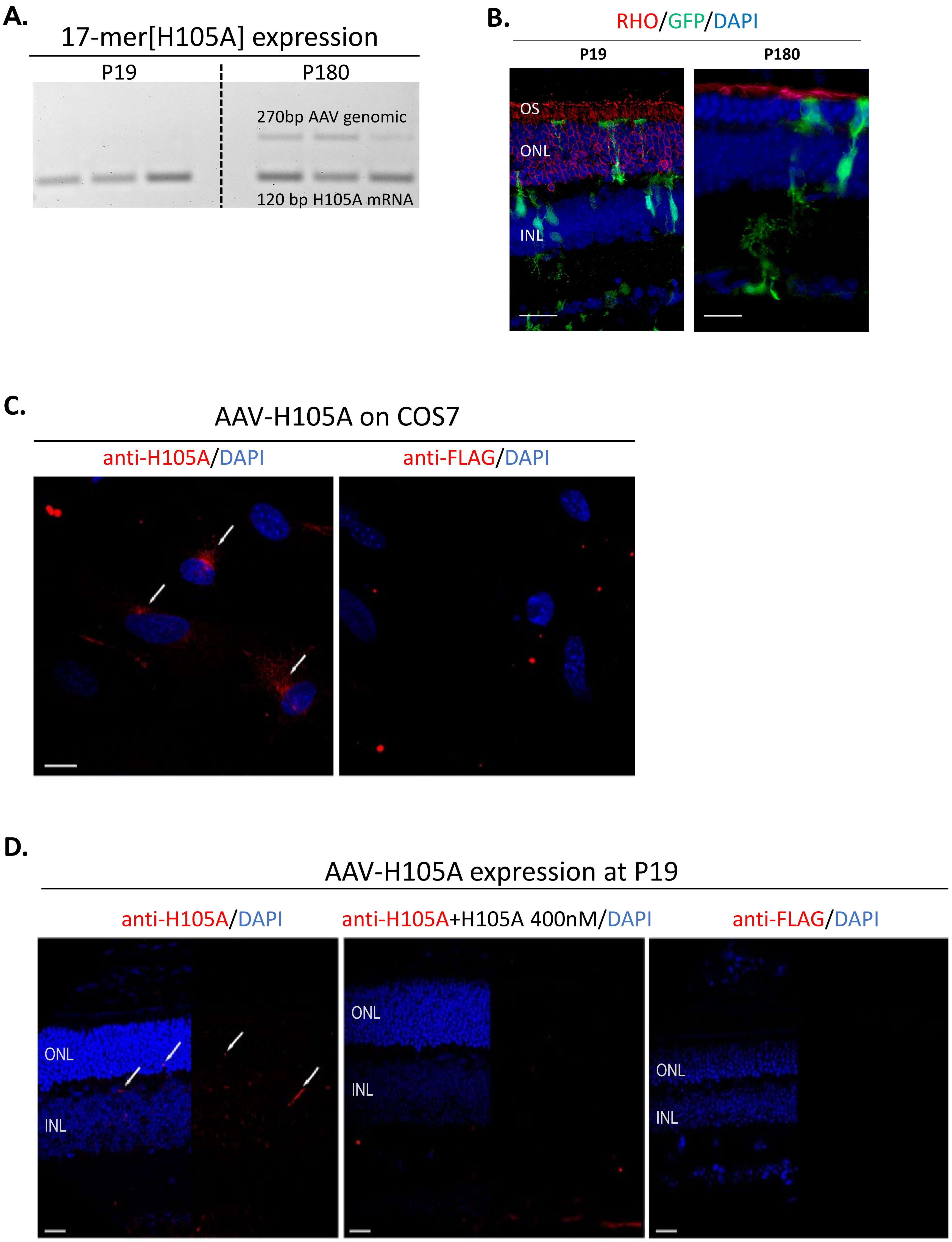

A.
17-mer[H105A] expression
P19
P180
270bp AAV genomic
120 bp H105A mRNA
B.
RHO/GFP/DAPI
P19
P180
OS
ONL
INL
C.
AAV-H105A on COS7
anti-H105A/DAPI
anti-FLAG/DAPI
D.
AAV-H105A expression at P19
anti-H105A/DAPI
anti-H105A+H105A 400nM/DAPI
anti-FLAG/DAPI
ONL
INL
ONL
INL
ONL
INL

**Figure S4.**

A.

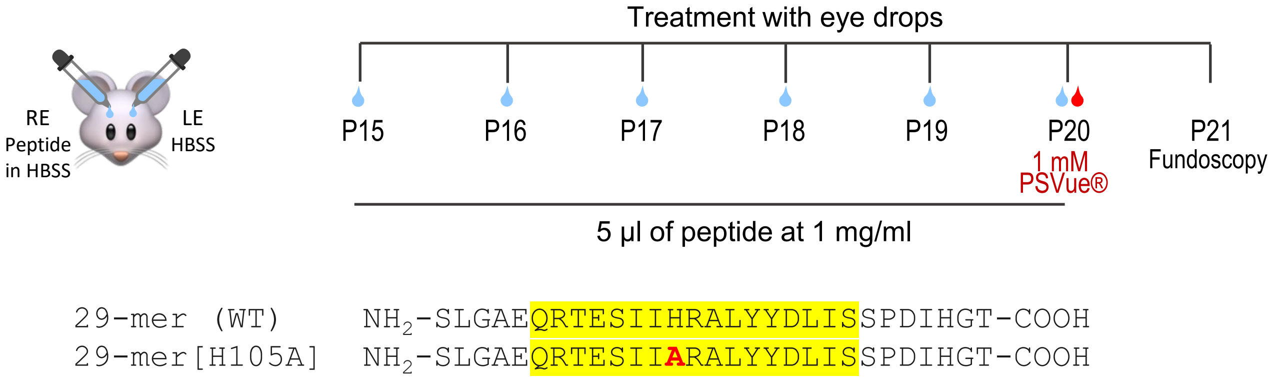


B.

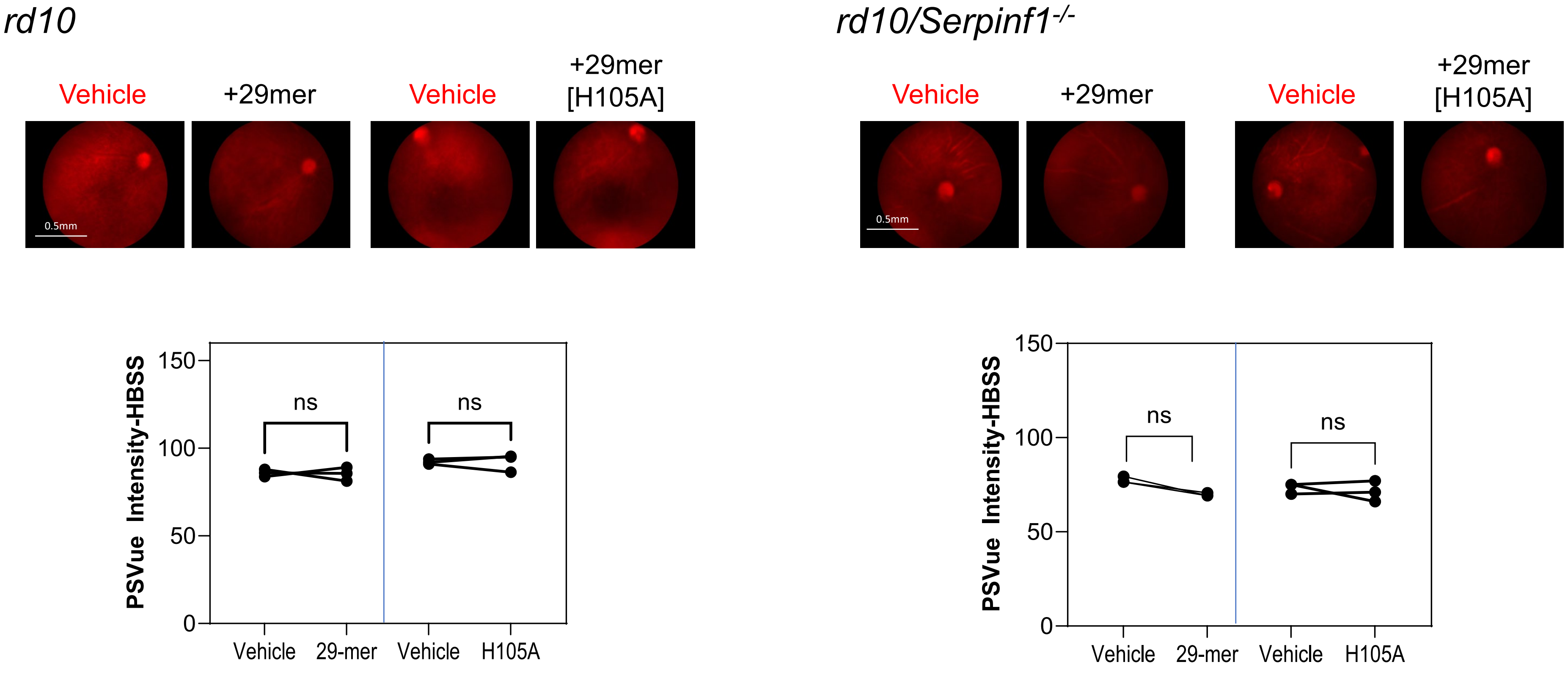


C. Alexa 488-labeled peptide did not penetrate in C57Bl/6J mouse eyes

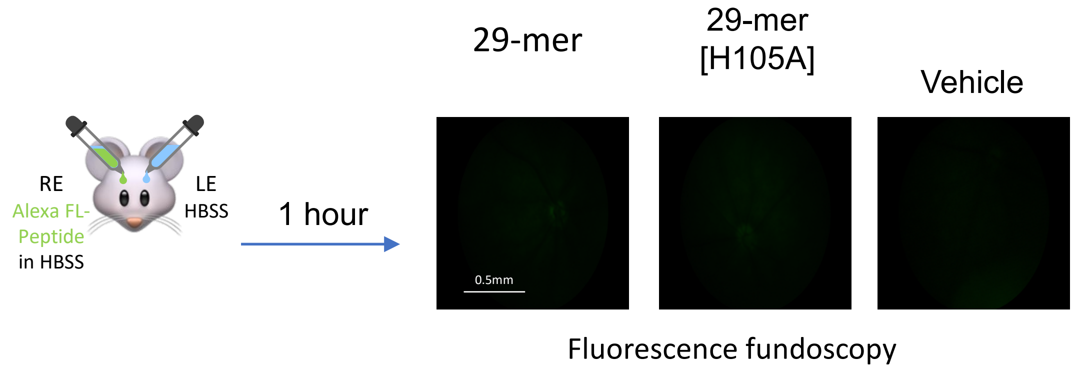

**Figure S5**.

A. Standard curve of AlexaFl488-17-mer, H105A and R99A peptides in mouse retinal extracts

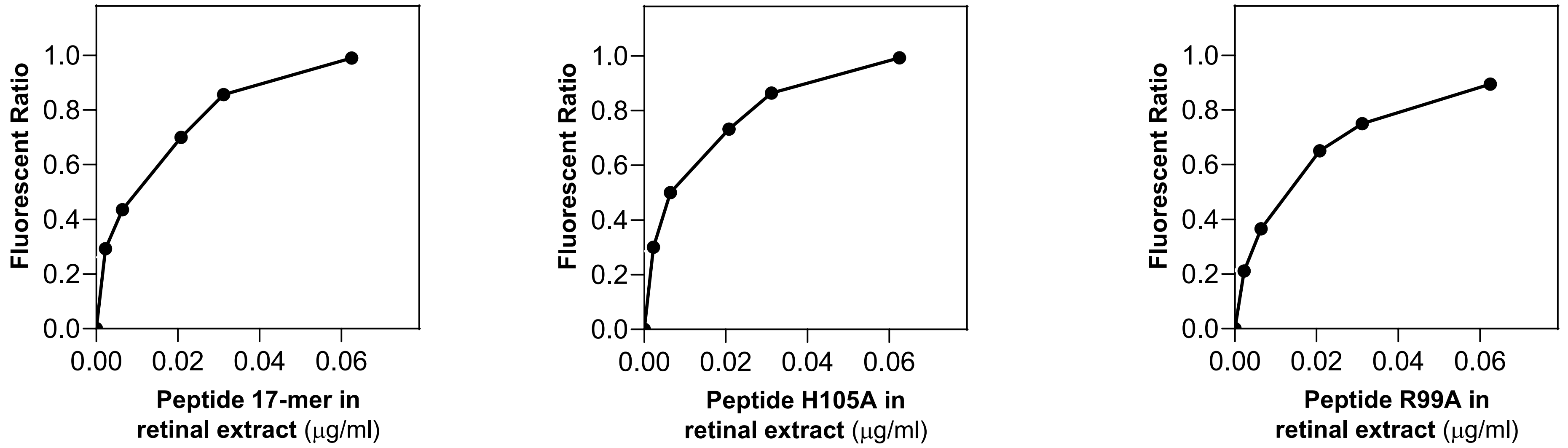


B. Effect of daily eyedrops of labeled PEDF peptides on ERG a-wave and b-wave retinal function

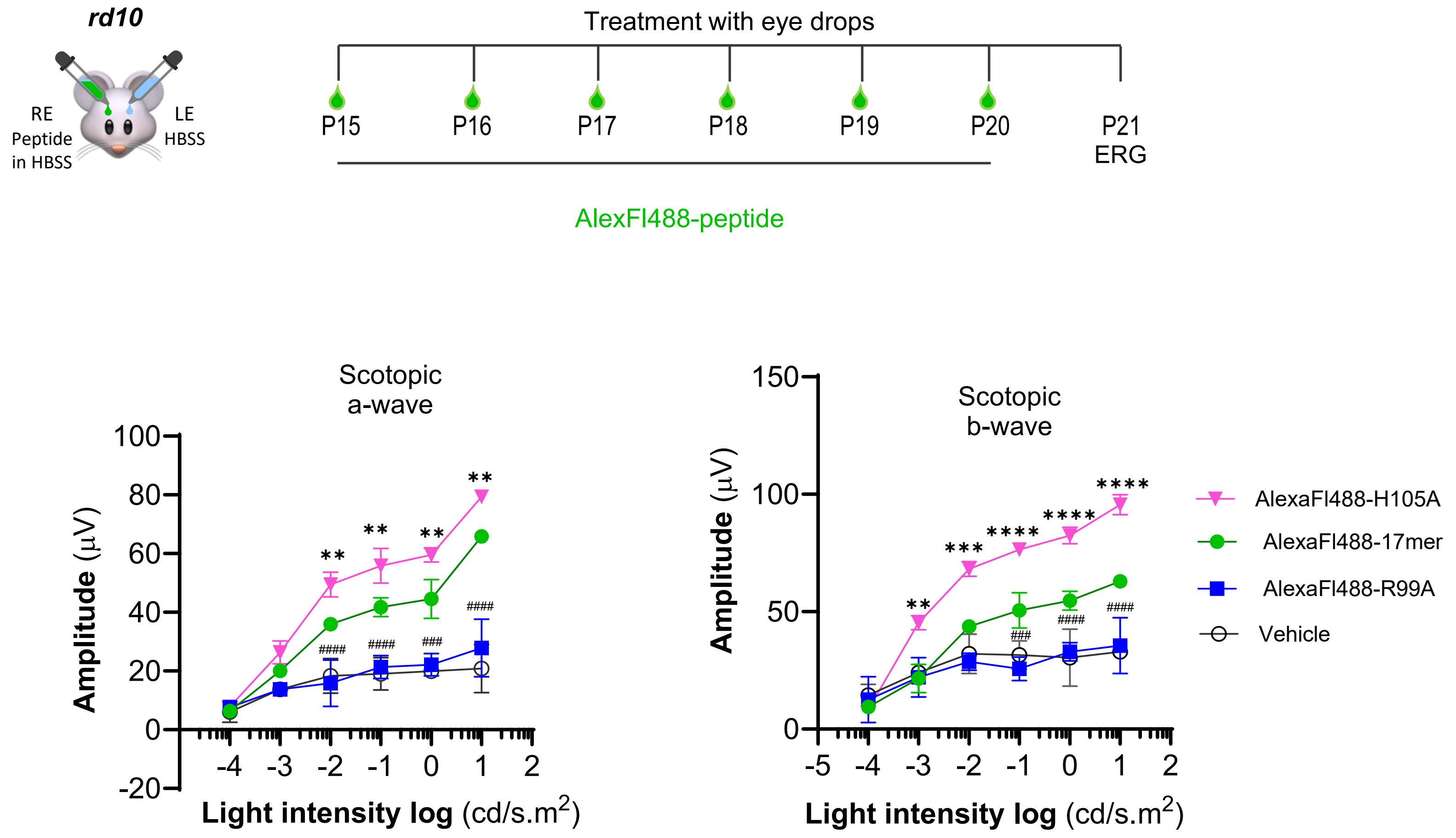

### SUPPLEMENTARY INFORMATION

**Figure S1. *In vivo* detection of dying photoreceptors in *rd10* mouse models by PS externalization monitoring.**

The highest PS externalization in fundi was at P17 for both models, being higher for *rd10* at all ages (**Fig. S1B**). The fluorescence slightly decreased at P21, to further decrease at P23 and being undetectable at P25. The fluorescence in mutant animals treated with HBSS or in wild type C57Bl/6J mice was undetectable and considered as background controls. To correlate these observations with photoreceptor viability, we compared the height of the outer nuclear layer (ONL) of photoreceptor cells in the retina. The ONL of mice at P15 and P21 showed nuclei that likely corresponded to photoreceptor cells undergoing early apoptosis before their near or complete loss by P25 (**Fig. S1C**). Depletion of photoreceptors at P25 explained the decrease in PSVue® fluorescence in fundi (shown in **Fig. S1B**) and precluded use of PSVue® to detect dying cells at P25 for both models. The data agree with previous reports in which in the *rd10* and *rd10/Serpinf1*$^{-/-}$ mice, loss of photoreceptor phosphodiesterase activity leads to photoreceptor degeneration, starting at postnatal day (P)16, and with near complete loss of photoreceptors by P25 and beyond *(13)*.

Given that cell death is largely regulated by the BCL2 family of interacting proteins, such as BCL2-Associated X protein (BAX) that plays a central role in apoptosis and B-cell lymphoma 2 (BCL2) as an anti-apoptosis regulator, we evaluated their distribution in photoreceptors of our mutant mice at P21. Immunofluorescence of retinal cross sections was performed to detect BAX and BCL2. The photoreceptors of both models exhibited BAX, while BCL2 was undetectable (**Fig. S1D**). Thus, PSVue® fluorescence fundoscopy of mice agreed with detection of photoreceptors undergoing early apoptosis in *rd10* and *rd10/Serpinf1*$^{-/-}$ mice in vivo, highlighting the age of P21 as endpoint for the rest of the study.

**(A)** Scheme showing the application of PSVue® 550 delivered via eye drops to *rd10* and *rd10*/*Serpinf*1 *null* mice at P16, P20, P22 or P24. Eye drops of 5 µl of a solution of 1 mM PSVue® in HBSS were applied per eye and fluorescence fundoscopy was performed in live animals 24 hours after application.

**(B)** *In vivo* detection of PS externalization in photoreceptors using PSVue®. Fluorescence fundoscopy micrographs of retinas at P17, P21, P23 and P25 of *rd10, rd10/Serpinf*1$^{-/-}$, and C57BL/6J mice after 24 h of administering PSVue® are shown. The right image on each row corresponds to retinas with HBSS (vehicle) only at P21, corresponding to background. Quantification of fluorescence intensity was done using ImageJ on images acquired by subtracting the background. Plots were generated using GraphPad. Each data point corresponds to the average of 3 images per retina for a total of 5 retinas per age.

**(C)** Retinal cross sections stained with DAPI showing the distribution of nuclei in *rd10* and *rd10/Serpinf*1$^{-/-}$ mice between P15 and P25. Representative images are shown. The dotted lines demarcate the outer nuclear layer (ONL) for photoreceptor nuclei.

**(D)** Retinal cross sections showing the distribution of BAX and BCL2 in *rd10* and *rd10/Serpinf*1$^{-/-}$ mice at P21 by immunofluorescence. Representative images are shown. For all immunofluorescence images shown, 5 retinas were used per group. ONL, outer nuclear layer, OS, outer segments, and IS, inner segments.

**Figure S2. Effects of extending the timeline or administering eye drops of 17-mer[H105A] peptide every other day in *rd10* mice.**

Eye drops of H105A peptide solution were administered to *rd10* mice either daily (**Fig. S2A**) or every other day (**Fig. S2D**). A total of 5 µl of a solution containing H105A peptide at 1 mg/ml in HBSS were applied as eye drops. The vehicle HBSS was added as eye drops in the contralateral eyes. When eye drops were added daily starting at P15 until P24, we noticed a rescue in the ONL thickness in *rd10* mice, as shown in **Figure S2B.** Eye drops also improved rod photoreceptor function in ERG in response to low intensity light at P25 (**Fig. S2C**). When eye drops of H105A were added every other day, a rescue in the ONL thickness in *rd10* mice compared to vehicle was also observed (**Fig. S2E**), with an improvement in a-wave of ERG, but no significant effect on the b-wave at P25 (**Fig. S2F**). The data demonstrate the efficacy of peptide H105A eye drops either daily or every other day in an extended timeline to stabilize photoreceptor morphology and light response function loss in *rd10* mice, however, these were less efficacious than daily H105A peptide eye drops.

**(A)** Scheme showing a timeline of peptide administration via daily eye drops of H105A peptide to *rd10* mice between P15 and P24. Assays were performed at P25.

**(B)** Representative microphotographs of retinal cross sections of *rd10* treated as in **Fig. S2A** and stained with hematoxylin and eosin are shown. To the right, a spider plot illustrating the thickness of the ONL of *rd10* mice treated with or without H105A as a function of distance from the optic nerve (ON). For all histology shown, 5 retinas per group were evaluated and each data point corresponds to the average ± SD per location relative to the ON per genotype by unpaired t-test. *$p < 0.05$, **$p < 0.001$, ***$p < 0.0001$, ****$p < 0.00001$.

**(C)** ERG of *rd10* mice at P25 and treated as in **Fig. S2A** were performed. Plots show amplitude (*y*-axis) for a- and b-wave as function of light intensity (cd/s.m$^2$, *x*-axis). For ERG data shown, the number of mice evaluated were $n$ =5 for *rd10* and each data point corresponds to the average ± SD of each genotype by unpaired t-test. *$p < 0.05$, **$p < 0.01$, ***$p < 0.001$, ****$p < 0.0001$.

**(D)** Scheme showing a timeline of peptide administration via eye drops of H105A peptide to *rd10* mice starting at P15 every other day until P23. Assays were at P25.

**(E)** Representative microphotographs of retinal cross sections of *rd10* treated as in **Fig. S2D** and stained with hematoxylin and eosin are shown. To the right, a spider plot illustrating the thickness of the ONL of *rd10* mice treated with or without H105A as a function of distance from the optic nerve (ON). For all histology shown, 5 retinas per group were evaluated and each data point corresponds to the average ± SD per location relative to the ON per genotype by unpaired t-test. *$p < 0.05$, **$p < 0.001$, ***$p < 0.0001$, ****$p < 0.00001$.

**(F)** ERG of *rd10* mice at P25 and treated as in **Fig. S2D** were performed in response to a STR. Plots show amplitude (*y*-axis) for a- and b-wave as function of light intensity (cd/s.m$^2$, *x*-axis). For ERG data shown, the number of mice evaluated were $n$ =5 for *rd10* and each data point corresponds to the average ± SD of each genotype by unpaired t-test. *$p < 0.05$, **$p < 0.01$, ***$p < 0.001$, ****$p < 0.0001$.

**Figure S3. H105A expression delivered by AAV2 is maintained at least for 6 months in the transduced retinas.**

As from the scheme in figure 5A, each *Rho*$^{P23H/+}$ mouse at P5 received AAV-H105A IVT in one eye and AAV-GFP IVT, a GFP expressing control AAV, in the contralateral eye. At P19 and P180, the mRNA and protein were assessed. **Figure S4A** shows a photograph of the gel with RT-PCR amplicons of H105A mRNA (120 bp) from transduced retinas, confirming the mRNA expression of the virus at P19 and P180. Fragments migrating as 270 bp correspond to AAV-H105A genomic DNA. Protein was assessed in retinal cross sections by immunofluorescence. **Figure S4B** shows mouse retinal cross sections with the fluorescence of the expressed GFP in green, DAPI in blue, and immunostaining of rhodopsin, a photoreceptor marker, in red, confirming that the AAV-GFP IVT primarily targeted ganglion cells and Müller glia cells. We also used a custom-generated anti-H105A antibody which was labelled with a FLAG-tag, and we validated the specificity of the antibody by several analyses in COS7 cells transduced with AAV-H105A and in sections of AAV-H105A transduced retinas. **Figure S4C** shows COS7 cells transduced with AAV-H105A and stained with anti-H105A antibodies and DAPI. Transduction of AAV-H105A in COS7 cells resulted in production of the neurotrophic factors as attested by the specific staining of anti-H105A (white arrows) in the cells and not detected with anti-FLAG and Alexa Fluor 568 anti-mouse secondary and tertiary antibodies, respectively (control in the absence of primary anti-H105A antibody). **Figure S4D** shows that in *Rho*$^{P23H/+}$ mouse retinas, AAV2-H105A IVT also resulted in production of H105A, as demonstrated by the specific staining of the anti-H105A (white arrows), that was blocked by competition with an excess of peptide antigen (H105A at 400 nM) and lack of staining with anti-FLAG and Alexa Fluor 568 anti-mouse secondary and tertiary antibodies, respectively. We concluded that a single IVT injection of AAV2-H105A in the *Rho*$^{P23H/+}$ mouse produced the neuroprotective agent in the retinal environment at P19 and P180.

**Figure S4. 29-mer and 29-mer[H105A] were not efficacious in protecting against photoreceptor cell death.**

A PEDF-derived peptide 29-mer (positions 93-121 of human PEDF) delivered by eye drops can prevent the development of experimental dry eye that affects the cornea *(56)*. We designed two peptides 29-mer and 29-mer[H105A] with the center of their sequences containing the 17-mer and H105A peptide sequences (in yellow highlight), respectively (**Fig. S5A**). To evaluate their penetration into the retina and therapeutic effect on mouse photoreceptor cell death, daily eye drops of 5 µl of 29-mer or 29-mer[H105A] at 1 mg/ml in HBSS were administered to left eyes and of HBSS (vehicle) to their contralateral right eyes of *rd10* and *rd10/ Serpinf1*$^{-/-}$ mice starting at P15 (**Fig. S5A**). At P21 PSVue 550 fluorescence fundoscopy showed no rescue in photoreceptor cell death with either peptide relative to vehicle in *rd10* and *rd10/Serpinf1*$^{-/-}$ mice (**Figs. S5B and S5C**). To evaluate the retinal penetration of 29-mer and 29-mer[H105A] peptides delivered via eye drops, we designed chemically synthesized peptides conjugated with Alexa Fluor™ 488 dye (Alexa 488) at their amino termini. Solutions of labeled peptides were administered to the left eye of C57Bl/6J mice and HBSS to the right eye. At 1-hour post-delivery, fluorescence fundoscopy showed no fluorescence intensity in the posterior retina for both peptides being similar to the fundi of the eyes that did not received peptide **(Fig. S5D)**. These observations indicated that both 29-mer and 29-mer[H105A] delivered by eye drops were unable to penetrate to the retina and not efficacious in protecting photoreceptors of *rd10* mice.

**(A)** Scheme showing a timeline of daily peptide administration via eye drops to *rd10* and *rd10*/*Serpinf*1 null mice between P15 and P21. The sequences of the peptides are given, and the yellow highlight corresponds to the 17-mer and 17-mer[H105A] region in the 29-mer and 29-mer[H105A], respectively.

**(B)** Mice were treated with peptides 29-mer and 29-mer[H105A] as indicated in **Fig. S5A**. Representative fluorescence fundoscopy micrographs of *rd10* and *rd10/Serpinf*1$^{-/-}$ retinas exposed to PSVue® are shown. Quantification of fluorescence intensity was performed using ImageJ of images acquired by subtracting the background of images of mice treated HBSS without PSVue®. Plots were generated using GraphPad. Each data point corresponds to the average of 3 ROIs per retina for a total of n = 3 retinas per genotype. Scale bar indicates 0.5 mm.

**(C)** AlexaFluor488-labeled 29-mer and AlexaFluor488-29-mer[H105A] were administered to C57Bl/6J mice at P21 days of age via eye drops (5 µl of a solution of each peptide at 1 mg/ml in HBSS per eye). Micrographs of fluorescence fundoscopy performed *in vivo* of the posterior retinas are shown. Scale bar indicates 0.5 mm.

**Figure S5. Alexa Fluor 488 labeled 17-mer, H105A and R99A peptides.**

(A) **Standard curves of Alexa Flour-488 labeled peptides in retinal extracts.** Plots of fluorescence of each labeled peptide in retinal extracts from C57Bl/6J mouse eyes are shown. Solutions of labeled 17-mer, H105A and R99A peptides at concentrations that ranged between 0.0 μg/ml and 0.0625 μg/ml were prepared. A total volume of 30 μl of each concentration was added to a well. Each data point corresponds to the average of two measurements per concentration.

(B) **Effect of daily eyedrops of Alexa Fluor-488 labeled peptides on ERG $_{a}$wave and $_{b}$wave retinal function of *rd10* mice.** ERGs were performed in *rd10* mice treated with eye drops of AlexaFl-488-17-mer, AlexaFl-488-H105A and AlexaFl-488-R99A, as shown in the scheme. Plots show amplitude (*y*-axis) as function of light intensity (cd/s.m$^2$, *x*-axis). For ERG data shown, the number of mice evaluated were $n$ =3 per condition and each data point corresponds to the average ± SD by unpaired t-test. ### $p < 0.001$ and #### $p < 0.0001$ for statistics between AlexaFl488-H105A and vehicle, and ** $p < 0.01$ and **** $p < 0.0001$ for statistics between AlexaFl488-H105A and AlexaFl488-17-mer. We note that eye drops of the labeled peptides AlexaFl-488-17-mer and AlexaFl-488-H105A improved retinal function as determined by ERG, but the labeled AlexaFl-488-R99A did not, as their unlabeled counterparts.

**Table S1. Peptides used in the study.**

| Peptide | Sequence |
|---|---|
| 17-mer | $NH_2$-QRTESIIHRALYYDLIS-COOH |
| H105A | $NH_2$-QRTESII**A**RALYYDLIS-COOH |
| R99A | $NH_2$-Q**A**TESIIHRALYYDLIS-COOH |
| 29-mer | $NH_2$-SLGAEQRTESIIHRALYYDLISSPDIHGT-COOH |
| 29-mer[H105A] | $NH_2$-SLGAEQRTESII**A**RALYYDLISSPDIHGT-COOH |
| AlexaFl-488-17-mer | AlexaFl-QRTESIIHRALYYDLIS-COOH |
| AlexaFl-488-H105A | AlexaFl-QRTESII**A**RALYYDLIS-COOH |
| AlexaFl-488-R99A | AlexaFl-Q**A**TESIIHRALYYDLIS-COOH |
| AlexaFl-488-29-mer | AlexaFl-SLGAEQRTESIIHRALYYDLISSPDIHGT-COOH |
| AlexaFl-488-29-mer[H105A] | AlexaFl-SLGAEQRTESII**A**RALYYDLISSPDIHGT-COOH |

**Table S2. Stability of 17-mer[H105A] peptide.**

| | Concentration (mg/mL) Day1 | Store | Concentration (mg/mL) Day8 |
|---|---|---|---|
| 1 | 1.04 | RT | 1.08 |
| 2 | 1.04 | 4 °C | 1.16 |
| 3 | 1.04 | -20 °C | 1.08 |
| 4 | 1.04 | -20 °C [FT]* | 1.28 |

*FT, Freeze and thaw cycles daily

**Table S3**. Antibodies used in the study.

| Antibody | Type & host | Application | Dilution | Company, catalog number |
|---|---|---|---|---|
| Anti-Bcl-2 | Rabbit Polyclonal | IF | 1:100 | Abcam, ab196495 |
| Anti-Bax | Rabbit Monoclonal | IF | 1:200 | Cell signaling, 14796 |
| Alexa Fluor 488 | Goat anti-Rabbit IgG (H+L) | IF | 1:200 | Thermo Fisher Scientific, A-32731 |
| Anti-RHO 1D4 | Mouse Monoclonal | IF | 1:500 | Santa Cruz, sc-57432 |
| Anti-GFP | Rabbit Polyclonal | IF | 1:200 | Thermo Fisher Scientific, A11122 |
| Anti-H105A | HuCAL, Monoclonal | IF, DB | 1:90 | BioRad, custom |
| Anti-FLAG | Mouse Monoclonal | IF | 1:100 | Merck, F3165 |
| Alexa Fluor 568 | Goat anti-Mouse IgG (H+L) | IF | 1:1000 | Thermo Fisher Scientific, A-11004 |

*IF, immunofluorescence; DB, dot blot
